# Multi-modal and multi-subject modular organization of human brain networks

**DOI:** 10.1101/2022.01.26.477897

**Authors:** Maria Grazia Puxeddu, Joshua Faskowitz, Olaf Sporns, Laura Astolfi, Richard F. Betzel

## Abstract

The human brain is a complex network of anatomically interconnected brain areas. Spontaneous neural activity is constrained by this architecture, giving rise to patterns of statistical dependencies between the activity of remote neural elements. The non-trivial relationship between structural and functional connectivity poses many unsolved challenges about cognition, disease, development, learning and aging. While numerous studies have focused on statistical relationships between edge weights in anatomical and functional networks, less is known about dependencies between their modules and communities. In this work, we investigate and characterize the relationship between anatomical and functional modular organization of the human brain, developing a novel multi-layer framework that expands the classical concept of multi-layer modularity optimization. By simultaneously mapping anatomical and functional networks estimated from different subjects into communities, this approach allows us to carry out a multi-subject and multi-modal analysis of the brain’s modular organization. Here, we investigate the relationship between anatomical and functional modules during resting state, finding unique and shared structures. The proposed framework constitutes a methodological advance in the context of multi-layer network analysis and paves the way to further investigate the relationship between structural and functional network organization in clinical cohorts, during cognitively demanding tasks, and in developmental or lifespan studies.

## INTRODUCTION

The human connectome is the complete set of white-matter connections among neural populations [52, 93]. This wiring diagram can be mapped and reconstructed using non-invasive imaging and represented as a graph or network [6]. The connectome shapes pattern of neural activity, inducing correlations - functional connections - between remote neural elements [37–39]. Human cognition and behavior are thought to be mediated by these distributed patterns of anatomical and functional connectivity among different brain areas [66, 74].

The relationships between structural and functional connectivity is a central concept in network neuroscience, and linking brain function to its architecture is a long-standing goal in brain research [57, 82]. A growing body of literature addresses this issue by trying to predict features of functional connectivity from structural features [71, 89] (for a review see [8, 97] or [5]). This has been done implementing statistical [76, 105] or communication [23, 48, 75] frameworks, considering the coupling with biophysical dynamic systems [1, 19, 27, 56, 96], or observing how brain lesions affect functional organization or link to behavior [98].

In addition to predicting one from another, structural and functional networks can be jointly analyzed to investigate common organizational properties, like modular structure, and see how they relate each other. Modular organization is a hallmark of brain networks [72, 73, 94] where groups of nodes are arranged into densely connected in communities that support specialized cognitive function [68]. This property has been observed in both anatomical and functional networks at different spatiotemporal scales [11]. In fact, brain networks can be partitioned into communities in many different ways, resulting in either small modules, made of functionally specialized areas [88], or large modules, hypothesized to support complex cognitive functions. A first contextual study [13] mapped structural communities (defined in terms of random walks) to patterns of functional connectivity, suggesting that such communities model brain function. More recently, in [40], the overlap between structural and functional weights and modules has been investigated during different states of integration and segregation of time-varying functional networks, finding that structure and function are closer when functional connectivity presents an integrated network topology. However, a straightforward analysis of the interplay between anatomical and functional modules is missing.

In parallel, a growing number of studies have begun investigating multi-layer network models of brain [25, 102]. The multi-layer framework allows for multiple instances or observations of a networked system to be analyzed under a single model. In the context of brain network analysis, multi-layer network models can be constructed to capture the covariance structure of functional brain data [7, 15, 18, 26, 85, 92] or to establish node correspondence across networks representing different subjects’ brains [17].

In this work, we introduce a novel extension of the multi-layer framework to directly investigate the relationship between anatomical and functional modular organization, which also accounts for the multi-scale nature of modules and their subject specificity. This approach is conceived as an extension of the well-known, and widely employed, multi-layer modularity maximization model [77]. Building on this work, we reformulate multi-layer modularity maximization by incorporating a new resolution parameter that effectively regulates the coupling between structural and functional connectivity matrices. Thus, we simultaneously map communities across subjects and modes of connectivity, which enables direct comparison of multi-modal modules via community labels. We applied this model to MRI-derived anatomical and functional brain networks of healthy adults, to investigate how the modular structure of brain networks varies across subjects and connectivity modality. We describe which brain sub-systems form modules consistent across modalities and those that decouple from one another to form modality specific modules. In summary, our work extends the multi-layer modularity maximization framework and paves the way for future studies to investigate structure-function relationships in different contexts.

## METHODS

### Experimental datasets and data processing

We leveraged anatomical and functional connectivity data belonging to two independently acquired datasets. We describe these data below.

#### NKI dataset

First, we considered data from the Nathan Kline Institute – Rockland Sample project [80] (NKI-RS, http://fcon_1000.projects.nitrc.org/indi/enhanced/). Institutional Review Board approval was obtained for this project at the Nathan Kline Institute (#226781 and #239708) and at Montclair State University (#000983 A and #000983B) in accordance with relevant guidelines. All participants gave written informed consent or assent. The anonymized dataset is freely available at http://fcon_1000.projects.nitrc.org/indi/enhanced/neurodata.html. The NKI dataset consists of imaging data from a community sample of subjects across a large portion of the human lifespan. We focused our analyses to subjects within the age range of 20-40 years old, to concentrate on structure-function relationships without added influence of age-related changes. The data processing resulted in anatomical and functional networks of 123 subjects. Below, we report an overview of the processing pipelines. For further details see also [63], where these pipelines have been discussed extensively.

#### Anatomical brain networks

Both T1-weighted (T1w) and diffusion (dMRI) images were provided, collected with a 3T Siemens Magneton Tim Trio scanner, using a 12-channel head coil. T1w images were preprocessed with the FreeSurfer (http://surfer.nmr.mgh.harvard.edu/) recon-all pipeline to reconstruct the Shchaefer network parcellation, which renders 100 cortical nodes (*N*). dMRI images were denoised, corrected for motion and susceptibility distortion, and then aligned to the corresponding T1w. Probabilistic streamline tractography was run using Dipy [41]. We extracted the structural connectivity matrices normalizing the number of streamlines that connect each region of interest (ROI) of the network parcellation, by the geometric mean volume of the connected ROIs (regions of interest) [12, 32]. In this way, we obtained weighted anatomical connectivity matrices where weights represented the connection density between brain regions. We performed rigorous quality control on these matrices excluding subjects based on the following criteria. T1w images were filtered based on artifact, such as ringing or ghosting and for FreeSurfer reconstruction failure. T1w images were also excluded if the scan was marked as an outlier in three or more of following quality metric distributions: coefficient of joint variation, contrast-to-noise ratio, signal-to-noise ratio, Dietrich’s SNR, FBER, and EFC. dMRI images were filtered based on corrupt data and artifact on fitted fractional anisotropy maps. Furthermore, dMRI data were excluded if the scan was marked as an outlier in two or more of following quality metric distributions: temporal signal-to-noise ratio, mean voxel intensity outlier count, or max voxel intensity outlier count. Tractography was run on images that had both quality controlled T1w and dMRI images. Tractography results were filtered based on artifact, which include failure to resolve callosal, cingulum, and/or corticospinal streamlines or errors resulting in sparse streamline densities. Outliers were computed as 1.5 times the interquartile range (only in the adverse direction) of the measurement distribution, relative to the whole NKI dataset. As a last step we converted these streamline density matrices into structural correlation matrices, as in [2], by computing the Pearson’s correlation between each pair of rows. In this way, the edge weights were restricted to the interval [− 1, 1] and the structural matrices are comparable to the functional, which is desirable in light of the construction of the multi-modal multi-layer network.

#### Functional brain networks

Functional images were preprocessed using fMRIPrep 1.2.5 [30], a Nipype [47] based tool. FMRIPrep is based on a boilerplate distributed with the software covered by a “no rights reserved” (CCO) license. It mainly uses the packages Nilearn 0.5.0, ANTs 2.2.0, FreeSurfer 6.0.1, FSL 5.0.9, and AFNI v16.2.07.

Each T1w volume was corrected for INU (intensity non-uniformity) using N4BiasFieldCorrection [4,101] and skull-stripped using the antsBrainExtraction.sh workflow. Brain surfaces were reconstructed using recon-all from FreeSurfer [24] and the brain mask estimated previously was refined with a custom variation of the method to reconcile ANTs-derived and FreeSurfer-derived segmentations of the cortical gray-matter of Mindboggle [61]. Spatial normalization to the ICBM 152 Nonlinear Asymmetrical template version 2009c [35] was performed through nonlinear registration with the antsRegistration tool, using brain-extracted versions of both T1w volume and template. Brain tissue segmentation of cerebrospinal fluid (CSF), white-matter (WM) and gray-matter (GM) was performed on the brain-extracted T1w using FSL’s fast [109]. Functional data was slice time corrected using 3dTshift from AFNI and motion corrected using mcflirt [58]. This was followed by co-registration to the corresponding T1w using boundary-based registration [49] with 9 degrees of freedom, using bbregister (FreeSurfer v6.0.1). Motion correcting transformations, BOLD-to-T1w transformation and T1w-to-template (MNI) warp were concatenated and applied in a single step using antsApplyTransforms using Lanczos interpolation.

At the end of this workflow we obtain NIFTI files for each participant. These have been subjected to a image quality control made with fMRIPrep’s visual reports and MRIQC 0.15.1 [29]. Functional data were excluded if greater than 15% of the frames exceeded 0.5 mm frame-wise displacement [83], or marked as outliers in 3 or more of the following quality metric distributions: DVARS standard deviation, DVARS voxel-wise standard deviation, temporal signal-to-noise ratio, framewise displacement mean, AFNI’s outlier ratio, and AFNI’s quality index. Outliers were computed as 1.5 times the interquartile range (only in the adverse direction) of the measurement distribution, relative to the whole NKI dataset.

A functional parcellation was used to define 100 brain regions (Schaefer 100 [91]) on the cerebral cortex in the same space of the BOLD NIFTI. This parcellation is designed to optimize both local gradient and global similarity measures of the fMRI signal. The nodes are also associated with canonical system labels [99].

At this point, we removed some of the nuisance variables from the BOLD signal of each cortical node (like motion throughout the scan). To do that we band-pass filtered (0.008-0.08 Hz) the data, applied a confound regression and standardized using Nilearn’s signal.clean, that removes confounds orthogonally to the temporal filters [64]. The regression was performed as in [90], because this strategy has been shown to be relatively effective for reducing motion artifacts [83]. Downstream, we obtained residual mean BOLD time series for each parcel, from which we computed the Pearson correlation to obtain the correlation coefficient that will fill the functional connectivity matrix.

Once obtained the structural and functional connectivity networks, we executed a final visual inspection of the matrices, which resulted in excluding subjects with average values of functional connections too high (*>* 0.2), or entropy values of the same edges too low (*<* 2*bits*) with respect to the others. We also excluded subjects whose correlation’s values between functional and structural connectivity weights were too low with respect to the others (*r <* 0.1).

#### HCP dataset

We also analyzed data from the Human Connectome Project (HCP) [103], a consortium projected to construct a map of human brain circuits and their relationship to behavior in a large population of healthy adults. The study was approved by the Washington University Institutional Review Board and informed consent was obtained from all subjects. It comprises a large cohort of subjects (*>*1000), from which multiple imaging data were acquired (diffusion MRI, resting-fMRI, task-fMRI and MEG/EEG), together with behavioral and genetic data. Details of the image acquisition and minimal preprocessing can be found in [45]. We focused only on the 100 unrelated subjects. Of the 100 subjects, four subjects were excluded as motion outliers, based on recorded motion during both the fMRI and dMRI acquisitions. These subjects’s mean framewise displacement measurements jointly exceeded 1.5 times the interquartile range of the distribution for all 100 subjects. One subject failed dMRI preprocessing. This resulted in obtaining structural and functional networks of 95 subjects. The processing of the data was made with the same cortical parcellation used for NKI data (Schaefer100), which led to networks made of 100 nodes.

### Multi-modal and multi-subject modularity optimization

The main objective of this work was to study the relationship between functional and structural connectivity from the perspective of their shared and unique modular structure. For this sake, we recovered communities of the brain networks through community detection [36], employing an algorithm based on modularity optimization. Modularity (*Q*) [44, 78, 79] is a global quality function that estimates how strongly communities are internally connected with respect to a chance level, so that optimizing *Q* results in the detection of assortative communities, i.e. groups of nodes that are internally dense and externally sparse. *Q* is defined as follows:

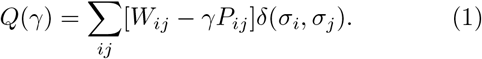

where *W*_*ij*_ and *P*_*ij*_ are the actual and expected weights of the link connecting nodes *i* and *j*. The variable *σ*_*i*_ ∈ {1, …, *K*} indicates to which cluster node *i* belongs, and δ(*x, y*) is equal to 1 if *x* = *y* and 0 otherwise. The parameter *γ* represents a spatial resolution weight that scales the influence of the null model, affecting the number and size of modules recovered. Using high *γ*-values results in many small communities and vice-versa.

A multi-layer version of *Q* has been introduced to detect communities in multi-layer networks [77], where layers correspond to estimates of the same network at different points in time, individuals, or connection modalities. The multi-layer analog of *Q* is defined as:

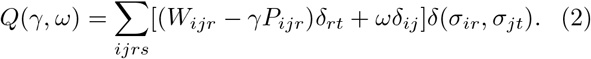

Nodes are linked to themselves across layers through the resolution parameter *ω* (Figure 1a). Its value affects the homogeneity of communities across layers (indicated through *r* and *t*), in a way that small *ω*-values emphasize layer-specific modular structure, while big *ω*-values point out communities shared across layers [87]. The optimization of *Q*(*γ, ω*) returns, for each layer, a partition of the network into assortative communities, whose dimensions and consistency across layers depend on two parameters of spatial and a cross-layers resolution. Applying multi-layer modularity optimization, instead of the single-layer one to each network, is generally convenient: the multi-layer approach is more resistant to noise [86] and returns community labels that are coherent across layers, so that one can easily track their evolution.

**FIG. 1.**
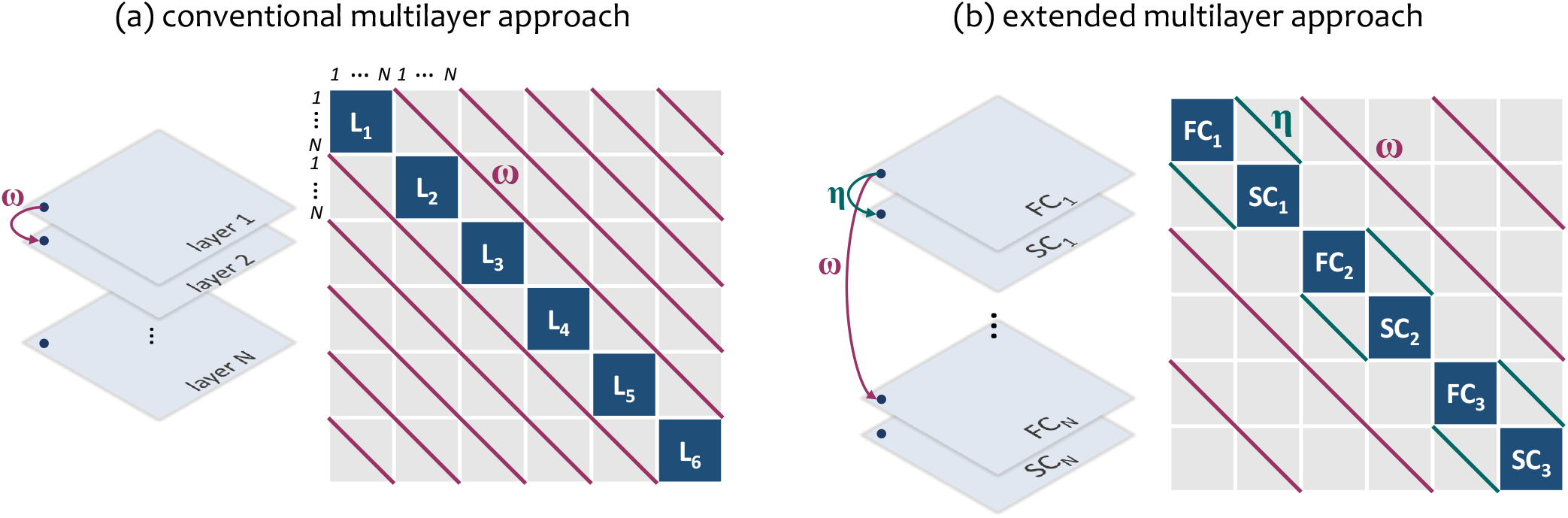
Schematic representation of the extended multilayer modularity optimization. a) conventional approach. Single-layer networks are aligned over the main diagonal of a square matrix of dimension [number of nodes × number of layers]. Modules are found by optimizing the modularity on this network in which each node is connected to itself across layers through the resolution parameter *ω*. b) extended approach. Functional and structural connectivity matrices of each subjects are aligned over the diagonal of a square matrix sized [number of nodes × number of subjects × 2 (type of connectivity)]. Modularity optimization is run on this network, considering each node connected to itself within modality and across subjects through *ω*, and between type of connectivity though *η*.

Historically, applications of this method have been, almost exclusively, to time-varying estimates of functional connectivity [7, 15, 18]. However, this multi-layer approach can be also used to link structural and functional modular organization. Multiple single-subject SC and FC matrices from the NKI can be averaged across subjects to form group representative networks. These, in turn, can be concatenated in a two-layers modularity matrix of dimension *N* ×*Modalities* (number od nodes × connection modality: 100 × 2) on which modularity optimization can be iterated with different combination of *γ* and *ω*. In this case, *γ* would impact the dimension of the communities, while *ω* their homogeneity between SC and FC.

In this work, we propose an extension of multi-layer modularity, formulated in order to encode information about different subjects and connection modality (SC or FC) simultaneously. Its expression is reported in Eq.3:

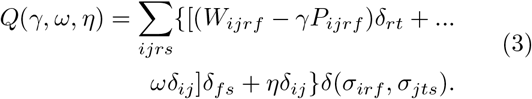

In addition to the spatial resolution parameter *γ*, we simultaneously consider two “temporal” resolution parameters, *ω* and *η*, that couple each node *i* with itself across subjects (indicated with *r* and *t*) and modalities (indicated with *f* and *s*) respectively (Figure 1b). In this way, *ω*, which we refer to as the *subject resolution parameter*, controls the homogeneity of the communities across all the participants, while *η*, the *modality resolution parameter*, regulates the coupling of the partitions between FC and SC networks within each participant. The *η* parameter operates in a way similar to *ω*, so that for increasing *η* values we obtain structural and functional partitions that are increasingly coupled. Thus, setting low *η* values would lead to partitions that highlight features that are unique to structure and function, while larger *η* values would correspond to features that are shared.

Thus, we built a multi-layer network as shown in Figure 1b, aligning along the main diagonal anatomical and functional modularity matrices of the 123 subjects. We obtained a square matrix of dimension *NS* × *N* × *Modalities* (number of subjects × number of nodes × connection modality: 123 × 100 × 2 = 24600). By optimizing *Q*(*γ, ω, η*) we partitioned nodes in single networks (the individual layers) into modules. We ran the modularity optimization varying the three resolution parameters in the range [*w*_*min*_, 1], with *w*_*min*_ = 2.1210^−6^ indicating the minimum weight of the SC matrices. In this range we considered 50 equally distanced values, so that we obtained an ensemble of 125000 partitions (50 × 50 × 50) for each subject and type of connectivity. We ran the optimization employing the openly-available genlouvain package (http://netwiki.amath.unc.edu/GenLouvain/GenLouvain), implemented in Matlab [59].

When running multi-layer modularity, the choice of the null model *P*_*ijrf*_ is crucial. There exist many possible definitions of null models, and we adopt the approach to set *P*_*ijrf*_ = 1 for all *i, j, s, f*. This is referred to as *uniform null model*, and it has been shown to be a good model to deal with correlation matrices [10, 100], resulting in communities with well-known topographic features. Since functional networks are rendered as temporal correlation matrices, and we converted anatomical networks into structural correlation matrices, we could use the uniform null model in all the network’s layers. As for the parameter *ω*, instead, it can be considered in two configurations: (i) all-to-all, meaning that nodes are linked to themselves through *ω* across all the layers (used if layers represent categorical variables); (ii) temporal, connecting nodes only between consecutive layers (used in time-varying networks). In this case, layers connected through *ω* represent different subjects, so that we used the first configuration.

### Analysis of the communities

#### Preliminary statistics on the communities

Through the multi-layer modularity maximization, we obtained a joint partition of anatomical and functional brain networks into nodal modules for each subject and for each combination of the resolution parameters. A widespread and easy approach is to focus on specific and fixed values of resolution parameters. Instead, we explored solutions over a range of parameters values, seconding the multi-scale organization of the brain networks [11]. Specifically, we assessed the modular structure by tuning the resolution parameters in the range [*w*_*min*_, 1] sampled with 50 values, obtaining 125, 000 partitions of the multi-layer networks for every subject’s SC and FC matrix. This corresponds to an ensemble of 250, 000 partitions for each subject (125, 000 ×2). Within this broad set of solutions, however, we restricted further investigation to a narrower subset, focusing on parameter combinations that result in neurophysiologically meaningful, non-trivial partitions. Criteria for selection included:

a. Number of clusters (*NC*). We avoid considering partitions with too many (or few) clusters comprised of too few (or many) nodes. As a range of interest, we focused on partitions of networks into 5-20 clusters, with an average of 20-5 nodes respectively (given that the network is made of *N* = 100 nodes). Hence, we only kept those partition solutions in which at least one subject is clustered more than 5 and less than 20 communities. Note that this criteria automatically handles singleton communities, i.e. communities comprised of just 1 node, by rejecting them.
b. Variability among subjects. We discarded partitions in which either functional or structural community structure was identical across all the subjects, as many studies demonstrate that the patterns of brain connectivity are subject-specific. We quantified the variability among subjects of node’s assignment through the normalized community en-tropy:

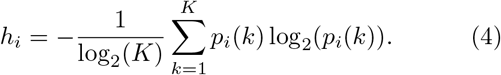

where *k* indicates the communities, and *p*_*i*_(*k*) is the fraction of subjects in which node *i* belong to community *k*. By dividing for log_2_(*K*) we normalized this measure in the range [0, 1], with 0 indicating identical assignments and 1 maximal variability. By averaging *h*_*i*_ for all the nodes, we obtained 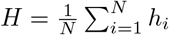, a measure of the variability of the entire partition across subjects, within each type of connectivity. We ignored resolution parameters providing *H* = 0 across subjects.

Based on these criteria, we considered *N*_*par*_ = 3732 combinations of {*γ, ω, η* } (out of the total 125, 000), in the ranges *γ* ∈ [0.02, 0.4], *ω* ∈ [2.1210^−6^, 0.16], *η* ∈ [2.1210^−6^, 1]. We based all the following analysis on this sample of partitions.

#### Modes of variability across modality and across subjects

In order to analyze the variability of communities across acquisition modalities, we computed the community entropy (Eq.5) between structural and functional partitions, which captures the variability of each node’s assignment. However, the precise entropy values depend on the specific combination of the resolution parameters {*γ, ω, η*}. Through Principal Component Analysis (PCA) we aimed to inspect the parameter space and see if there existed patterns of variability that are recurrent inside this space. Thus, for each point in the parameter space (identified by different combinations of {*γ, ω, η*}), we computed the entropy between partitions coming from functional and structural networks, and we averaged this value across subjects. We aligned the resulting vectors in a matrix *H*_*sf*_ of dimension *N* × *N*_*par*_, and we subjected it to Singular Value Decomposition (SVD). SVD decomposes the matrix *H*_*sf*_ into singular vectors *U*_*sf*_ ∈ [*N* × *N*] and *V*_*sf*_ ∈ [*N*_*par*_ × *N*], and singular values Σ_*sf*_ ∈ [*N*_*par*_ × *N*], so that:

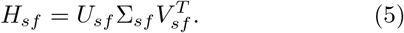

The matrices *U*_*sf*_ and *V*_*sf*_ are orthonormal by definition and contain the principal component scores and coefficients, respectively. The columns of *U*_*sf*_ can be interpreted as the modes of variability between SC and FC partitions, while the values in the rows in *V*_*sf*_ indicate where these modes are likely to appear in the parameter space. The diagonal elements of Σ_*sf*_ contain the information about the covariance with which each component explains the variability across SC and FC.

Limiting the study to the components explaining most of the variance between SC-FC community entropy, we identified the points in the parameter space where these components are most strongly expressed. In this way, we divided the parameter space into non-overlapping regions that correspond to distinct patterns of coupling between structural and functional modules. Within each one of these regions we extracted representative partitions for both structural and functional networks, by computing the Variation of Information (VI) [70], an information theoretic measure of distance between pairs of partitions, and recorded the centroid, the partition least distant from all other partitions (i.e. with lowest VI). A schematic representation of these analysis can be found in Supplementary Material, Figure S1 and S2.

We performed a similar analysis to identify modes of variability across subjects, focusing our analyses on inter-subject variation of anatomical or functional modules, independently. We built a matrix *H*_*sbj*_ of dimension *N* × *N*_*rep*_ containing in each column the vector of the entropy *h* computed among subjects within modality each modality (structural/functional connectivity). We executed the SVD to decompose *H*_*sbj*_ in the singular vectors *U*_*sbj*_ ∈ [*N* × *N*] and *V*_*sbj*_ ∈ [*N*_*par*_ × *N*], and singular values Σ_*sbj*_ ∈ [*N*_*par*_ × *N*]:

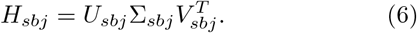

We did this for both structural and functional partitions. Again, the matrices *U*_*sbj*_ and *V*_*sbj*_ are orthonormal and contain the principal component scores and coefficients. Here, we interpret the columns of *U*_*sbj*_ as the modes of variability of the partitions across subjects. The values in the rows in *V*_*sbj*_ indicate where these modes are likely to appear in the parameters space, while the diagonal elements of Σ_*sbj*_ contain the variance with which each component explains the variability across subjects.

#### Assessment of the segregation and integration of the network’s modules

Next, we analyzed the structural and functional partitions in terms of single-layer modularity (Q) and participation coefficient (*pc*_*i*_) [51]. The participation coefficient (Eq. 7) quantifies the distribution of each node’s edges across modules. High *pc*_*i*_ values reveal nodes well integrated in the system, across communities, and vice versa.

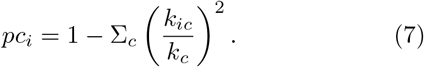

Where *k*_*ic*_ is the sum of the edges of node *i* within modules *c*, and *k*_*c*_ is the sum of the edges inside module *c*.

Through the single-layer *Q* (Eq.1) we assess how modular the partitions are by comparing the fraction of edges within each community to a null model. Together with the modularity score of the anatomical (*Q*_*s*_) and functional (*Q*_*f*_) partitions, we computed the structural-functional modularity score *Q*_*sf*_ [54]. *Q*_*sf*_ captures the modularity of the functional modules imposed on the anatomical networks, returning the level of overlap between the two. Finally, in order to take the global modularity score back to the topology of the brain networks, we inferred how single modules contribute to *Q*. This could be done from Eq.1 by summing the elements of the sub-matrix referring to a modules *K*.

## RESULTS

The brain modular structure reveals groups of brain regions that are functionally or anatomically related. Identifying this architecture at different scales can lead to important clinical and behavioral insights. In this work, we introduced an extension of the multi-layer modularity maximization, developed to analyze multi-subject multi-modal datasets. By concatenating structural and functional connectivity matrices of different subjects and treating them as coupled layers of a multi-layer network, our approach is meant to simultaneously map brain communities into subjects and type of connectivity at different scales. In the following sections we illustrate the results of the application of our new framework to anatomical and functional networks from the NKI dataset.

### Multi-modal modular structure with conventional multi-layer modularity optimization

As a first step, we show how topological properties of anatomical and functional networks can be linked through the conventional multi-layer modularity optimization framework (Figure 1a). To this end, we extracted representative anatomical and functional group-averaged networks for the NKI dataset by averaging SC and FC matrices across the 123 subjects. Hence, we built a two-layers modularity matrix and we iterated the conventional multi-layer modularity optimization (Eq.2) 100 times with combinations of *γ* and *ω* whose values span the range [0, 1]. Then, we retained only partitions with non-singleton communities, which led to consider *γ* in the range [0, 0.83]. Here, we comment how the two resolution parameters affect the optimization’s output in terms of coupling between anatomical and functional modules and how we can study their relationship at different scales.

By computing the cross-layer VI and nodal entropy (that are the VI and entropy between FC and SC partitions), and averaging across the 100 iterations, we show how the resolution parameter *ω* controls the homogeneity of modules across layers, and thus, in our case, connection modalities. These indices measure how much nodes’ assignments to a community vary at the node level (entropy) and globally (VI). As expected, entropy values decreased monotonically with *ω* (Figure 2(a)), which means that higher *ω*-values leads to high coupling between FC and SC partitions. Analogously, VI decreased monotonically with *ω* (Figure 2(g), blue line), meaning that also globally FC and SC partitions tend to get closer one another, until *ω >* 0.7, at which point the partitions are identical (*V I* = 0, *H*_*sf*_ = 0). Thus, by tuning *ω* in its range, we were able to explore different levels of structure-function relationships, highlighting modular configurations that are distinctive of the connection modality, or shared between them.

**FIG. 2.**
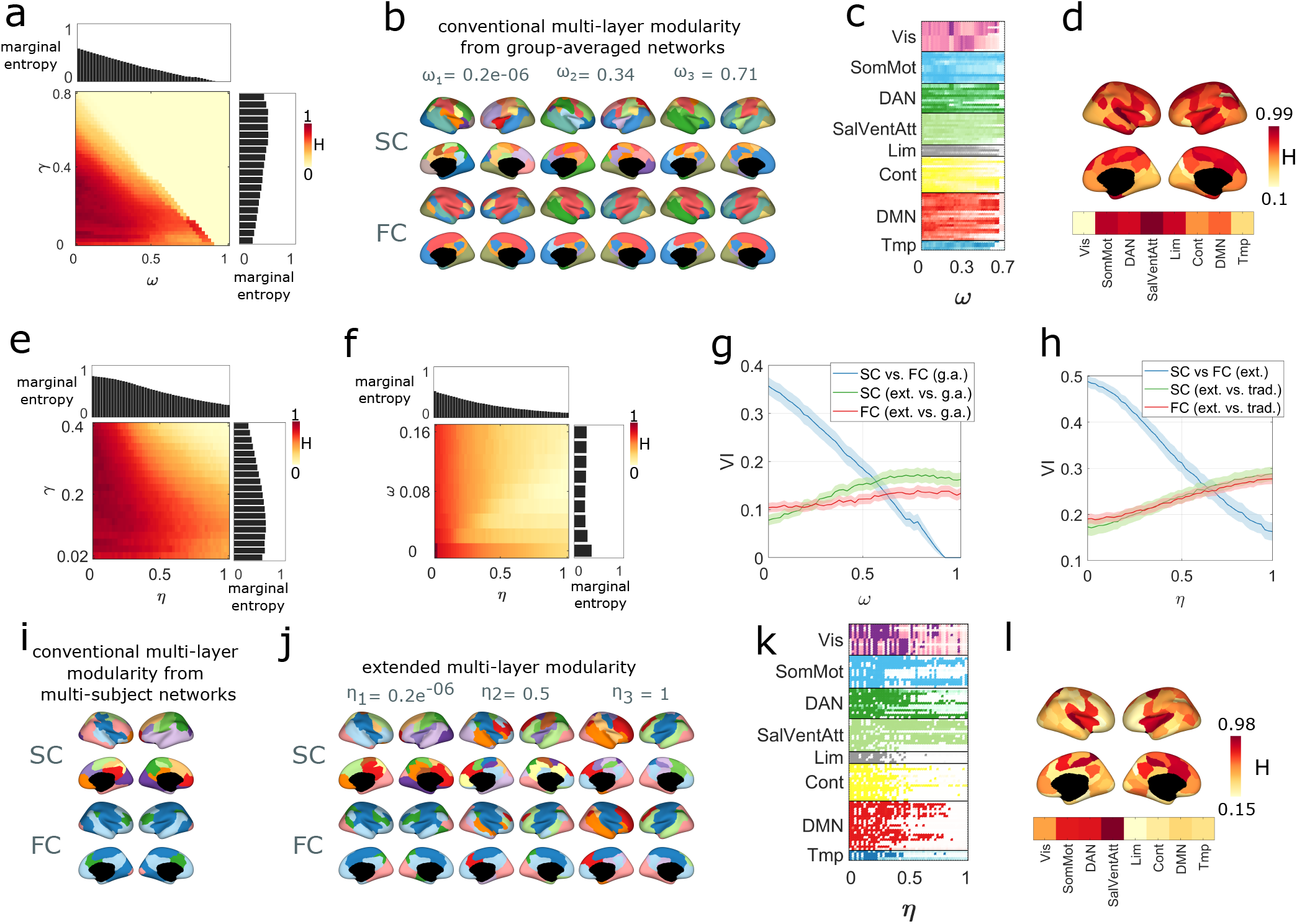
Multi-layer modularity to couple structural and functional communities. (a) Entropy between types of connectivity observed in the group-average analysis varying the resolution parameters *ω* and *γ*. (b) Projection on the brain cortex of the structural and functional partitions obtained with multi-layer modularity optimization with *γ* = 0.16 and increasing *ω* on the group-averaged networks. (c) Node’s entropy between SC and FC partitions in the group-average networks in the *ω*-range (*γ* = 0.16), and its average projected in the brain surface (d). (e) Entropy between types of connectivity in the extended model by varying the resolution parameters *η* and *γ* and having fixed *ω* (average across subjects). (f) Entropy between types of connectivity in the extended by varying the resolution parameters *η* and *ω* and having fixed *γ* (average across subjects). (g) Distance between SC and FC partitions in the group-average analysis across the *ω*-range (blue line). The red and green plots report the distance between the functional and structural partitions obtained with the extended model and those obtained in the group-average analysis. (h) Distance between SC and FC partitions in the extended model across the *η*-range (blue line) in terms of Variation of Information (mean and confidence interval). The red and green plots report the distance between the functional and structural partitions obtained with the extended model and those obtained applying conventional multi-layer modularity maximization on the SC and FC multi-subjects networks. (i) Projection on the brain cortex of the consensus partitions obtained applying the classic multi-layer modularity optimization on the multi-subject networks with *γ* = 0.12 and *ω* = 0.06. (j) Projection on the brain cortex of the consensus partitions obtained with the extended multi-layer modularity optimization optimization for increasing *η* (*γ* = 0.12; *ω* = 0.06). (k) Node-wise entropy between SC and FC partitions (averaged across subjects) in the *η*-range (*γ* = 0.12; *ω* = 0.06), and its average projected on the brain surface (l).

By definition, *γ* affects the size of the detected communities. However, in Figure 2(a), one can see how *γ*-values also impact the cross-layer homogeneity, in a proportional way (even if to lesser extent with respect to *ω*, as shown by the marginal plots). This is trivially explained by the progressive diminishing of the dimension of the modules. In fact, any two partitions with a number of clusters close to the number of nodes are intuitively more similar. Also the lowest *γ*-values brought to more coupled partitions, and for a similar reason (few bigger clusters are more likely to comprehend the same nodes). Further statistics and plots regarding the effect of the spatial resolution parameter on the dimension of the discovered communities have been reported in Supplementary Material, Figure S4.

Focusing on a specific *γ*-value (*γ* = 0.16), we reported in Figure 2(b) an example of how modules in SC and FC networks gradually reconfigure, increasing *ω*, to meet a higher overlap. The most evident change, by eye, happens in the areas covering the primary motor cortex. These belong to hemispheric-specific clusters in the SC partitions for low *ω* (as expected, since structural connectivity favors short-distance connections), but then increasing *ω* they start participating in the same modules, as in FC partitions. Unpacking the nodal entropy (Figure 2(c-d)) we could better observe these local properties and how different brain areas reconfigure at a different rate. For instance, we found that ventral attention network present high entropy until *ω* = 0.7 (where FC and SC partitions become identical), while visual and temporal areas participate in modules that are overlapped in SC and FC partitions even before *ω* = 0.7. Note that in Figure 2(b,c) we showed modular structure and nodal entropy up to *ω* = 0.7. We excluded the possibility of a perfect alignment between structural and functional modular organization, given the different nature of the signals and the fact that structural modules underpin a static wiring of anatomical pathways, while functional modules are shaped by dynamic statistical relationships between brain areas.

In summary, we showed how conventional multi-layer modularity optimization can be used to investigate the coupling between structural and functional topological properties. However, applied in this manner, this approach can only operate on averaged brain network data, ignoring single-subject’s information. Indeed, this model is able to track communities along only one dimension, which we can choose to be time (e.g. as in [7]), subjects (e.g. as in [17]), or connection modality (as shown here). In the next section we illustrate how the extension that we propose overcomes this limit. For sake of clarity, note that in the just presented conventional multi-layer framework the parameter *ω* couples SC and FC networks. In our extension instead, SC and FC will be linked by a new parameter *η*, while *ω* will bridge matrices across subjects. This is well explicated in Methods, Eq.2 and Eq.3.

### Detection of multi-subject multi-modal modular structure

In the previous section we illustrated how the multilayer framework could be applied to multi-modal network data, focusing on group-averaged networks. Here, we show how the extended multi-layer modularity optimization (Eq.3) can detect modules in structural and functional networks simultaneously from the 123 subject of the NKI dataset. This time, the output of the optimization algorithm depends on three resolution parameters {*γ, ω, η* }, that we varied in the range [*w*_*min*_, 1]. One challenge is that some of the parameters will yield uninformative partitions. We illustrate how we excluded unwanted partitions and, once restricted the space of investigation, how {*γ, ω, η* }-values affect the number of modules and their coupling across subjects and connection modality.

To retain only informative partitions, we studied how the spatial and cross-subject resolution parameters, *γ* and *ω*, impact the statistics of the communities and we restricted their range consequently. According to our formulation of the multi-layer framework, the spatial resolution parameter, *γ*, influences the number of communities, while *ω* impacts the homogeneity of the partitions across subjects. Here, we computed the optimization while tuning the parameters in the range [*w*_*min*_, 1], and we observed how the number of clusters and community entropy are distributed in the parameter space. Results are reported in the Supplementary Material, Figure S3. As expected, we found that setting *γ* to high (low) values leads to many small (few big) clusters, while increasing *ω* leads to lower variability of the modular structure across subjects (Figure S3(f-i)). We selected non-singleton partitions with 5-20 communities, where nodes’ assignment to the communities differs for at least one subject (i.e. entropy within modality greater than 0). This procedure narrowed the *γ* and *ω* ranges down, into a non-rectangular parameter space limited by *γ* ∈ [0.02, 0.4], *ω* ∈ [2.1210^−6^, 0.16], where anatomical and functional partitions are respectively made of *NC*_*s*_ = 16 ± 9.8 and *NC*_*f*_ = 8 ±4.8 clusters. Each combination of the parameters in this subspace corresponds to an informative partition. As a demonstration, we computed the distance of all the partitions from a set of canonical brain systems [99], observing that the minimum distance falls in the selected subspace (see Supplementary Material, Figure S5).

Once we identified a sub-space of meaningful partitions through the parameters *γ* and *ω*, we could observe how the *η* parameter enables us to explore different levels of coupling between structural and functional modular structure. In our model *η* determines the extent to which identified communities persist across types of connectivity. The hypothesis is that there would be significant differences among structural and functional partitions when the coupling parameter is low, but the partitions would converge increasing *η*’s value. To test this hypothesis we computed the VI between structural and functional partitions for each *η*-value, averaging in the restricted {*γ, ω* } ranges. The results, reported in Figure 2(h), showed a decreasing trend of VI for increasing *η*-values (blue line). Low *η*-values yield weak coupling between FC and SC partitions, while high *η*-values yield strong coupling between the same partitions. This supports the hypothesis that the brain modular structure differs between types of connectivity, but that adjusting *η* we can encourage the algorithm to find features that are unique to function and structure, as well as shared features. Note that this is analogous to what we found using the connection modality as layer-dimension in the group-averaged conventional model (Figure 2(g)). The only difference is the range of VI-values, which is shifted down in the conventional case. This might be an effect of the group-average, that leads to flatten network’s properties over the sample.

Furthermore, in Figure 2(h), we showed the VI computed between the partitions obtained with the new model and those obtained applying classic multi-layer modularity optimization to the multi-subject matrix comprising either all SC or FC networks (green and red lines). In this case the VI tends to increase with *η*, meaning that the higher *η* is, the more that the multi-layer partitions diverge from the traditional ones. However, these VI-values remain low (*<* 0.3) along the *η*-range, so that increasing *η* affects the coupling between SC and FC partition to a greater extent in the new framework, than the divergence of such partitions with the traditional ones.

The parameters *γ* and *ω*, are responsible for modulating the number of clusters and homogeneity of the partitions across subjects (Figure S3). We showed that the coupling between structural and functional modules is analogously governed by *η*. A further demonstration of this is in Figure 2(e-f), where we reported the normalized nodal entropy (H) between SC and FC partitions, first calculated within subjects and then averaged across all subjects, in the restricted parameter space. As expected, we found that H varies monotonically only with *η*, decreasing when such parameter increases. Overall, we observed that H varies all over the parameter space and it shows its lowest values, meaning high coupling between FC and SC partitions, when all the parameters {*γ, ωη*} are set to high values.

Our multi-subject and multi-modal modularity is meant to detect modules of different sizes and more or less consistent across subjects and types of connectivity. However, in some cases, instead of observing how global properties of the modules vary across the whole parameter space, one may want to investigate a more focused set of partitions and provide a more detailed description of the community structure. Our algorithm allows for this possibility. For instance, in Figure 2(j-l), we isolated the structural and functional partitions obtained with *γ* = 0.12 (meaning on average *NC*_*s*_ = 13.9 ± 1.7 and *NC*_*f*_ = 7.3 ±1.9) and *ω* = 0.06 (ensuring good consistency across subjects). We observed how in this specific case the partitions become progressively coupled across *η* (Figure 2(j)). The, we also observed how much they differ compared to the ones obtained with the conventional approach where the modularity optimization is applied to the multi-subject matrix containing all FC (or SC) networks (Figure 2(i)). With low *η* the two approaches led to similar partitions (*V I*_*s*_ = 0.32; *V I*_*f*_ = 0.24). Nevertheless, increasing its value gradually modifies the output in order to detect common sub-systems. To further explore structure-function relationship, we can also look at which brain regions are more prone to participate in similar communities in SC and FC networks for increasing *η*-values, and which instead remain modality-specific (as we did in the previous section). In the subspace defined by {*γ* = 0.12, *ω* = 0.06} we observed that the DMN together with the temporal, limbic and control systems belong to the first category, while the somato-motor area, DAN and salience and ventral attention networks belong to the second one (Figure 2(l)). The same analysis can be used to explore communities at coarser or finer scales, meaning focusing on lower or higher *γ*-values (see Figure S6). Furthermore, in a similar way, one can also explore how either structural or functional modular organization reconfigures across subjects (see Figure S10).

Here, we have illustrated the community structure that can be obtained using multi-layer multi-modal modularity maximization. Its output depends on three resolution parameters whose value impacts modules size as well as variability across subjects and connection modality. Predefined heuristic about the output community structure allowed us to focus on parameter ranges where the character of detected communities is consistent with previous reports, while still capturing meaningful inter-subject and inter-modality variability. We also showed how our model allows for the coupling between structural and functional partitions to be systematically adjusted. A more detailed description of the modular structure in the parameter space will follow.

### Modes of variability between connection modalities

In the previous section we identified a subset of parameters where partitions could be considered meaningful for imaging-based analyses. Inside this subspace, we showed in which way the three resolution parameters {*γ, ω, η*} affect the scale of the partitions and the magnitude of their variability between types of connectivity and among sub-jects. Now, we want to discover if there exist recurrent patterns of inter-modality variability which are encoded in the parameters subspace in a non-random manner. In fact, knowing how the SC-FC coupling depends on the scales of the communities is crucial in more pragmatic study, for instance with clinical implications. For this purpose, we stored in a 100 × 3732 (*N* ×*N*_*par*_) matrix the vectors containing the normalized nodal entropy across modalities (one vector for each point in the sub-sampled parameter space) and we performed the Principal Component Analysis (PCA) on this matrix. Through the PCA we obtained 99 principal component scores (in the form of orthonormal vectors of length equal to the number of nodes), and the relative coefficients, informative of their contribution on each of the 3732 rows of the data. Thus, we identified the modes of variability projecting the scores and the coefficients on the cortex surface and on the parameter space, respectively.

In Figure 3 we reported the results relative to the first five components, that, out of the 99, explain most of the variance of the SC-FC community coupling (12.53%, 9.67%, 7.77%, 6.17%, 5.69% respectively), while the remaining components each explains less than 5% of the variance (see Supplementary Figure S7).

**FIG. 3.**
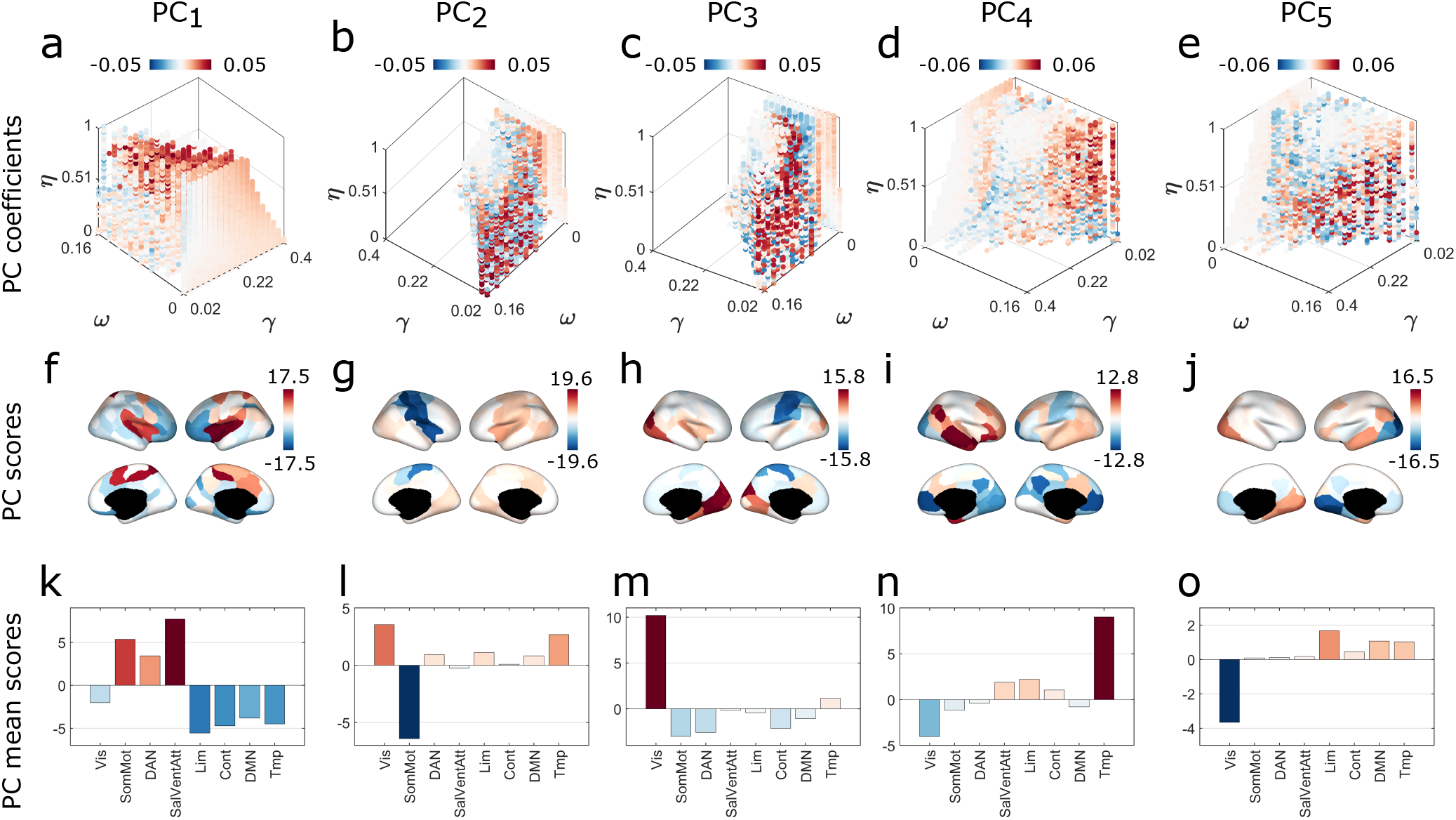
Principal component analysis (PCA) for the detection of modes of variability between structural and functional community structure. Columns of this figure correspond to the first five components of the decomposition. (a-e): projection of the first five PC coefficients into the parameter space identified by the parameters *γ, ω, η*. (f-j): projection on the cortex surface of the first five PC scores. (k-o): average within the functional systems of the first five PC scores. In each component, brain areas colored with red present community assignment highly variable between functional and structural networks. These variations mostly occur in the space of parameter identified by red dots. Blue-colored brain areas instead, identify stable node’s community assignment in the relative blue points of the parameter space.

Globally the first five *PC*_*s*_ occupied different zones in the three-dimensional parameter space (Figure 3(a-e)). *PC*_1_ was expressed at high *η* and *γ* values, corresponding to a subspace in which inter-modality variability was low and the brain is partitioned into fine modular structure, i.e. many small modules. Projecting *PC*_1_ scores onto the cortex showed that, in this region of the parameter space, the nodes whose module’s assignment differs most between FC and SC networks belong to the Dorsal Attention Network (DAN), the Salience and Ventral Attention Network and the somatosensory and motor areas (Figure 3(f,k)). On the contrary, other regions like those from the Control, Default Mode Network (DMN), Limbic, and Temporal systems, showed higher consistency between the two connectivity measures within this subspace. The other principal components were expressed in other regions of the parameters space and showed different patterns of inter-modality variability, suggesting that patterns of variability of the SC and FC modular structure were resolution-dependent. For example, *PC*_2_ coefficients assumed high positive values in the subspace where *γ* and *η* are low and *ω* is high, corresponding to networks parsed in few clusters, highly coupled across subjects and barely coupled between connection modality. Here, the regions whose module assignments were more variable between FC and SC networks are those involved in the visual and temporal systems. The regions belonging to the somatosensory and motor systems instead, show higher consistency between modalities. In *PC*_4_ coefficient are more expressed at low *γ*, and high *ω* and *η*, that is where partitions present few clusters, high correspondence among subjects and a good level of overlap between modalities. In this subspace the visual and somatomotor systems, together with the DAN and DMN, tend to maintain the same modules allegiance in SC and FC networks, while the regions mostly located in the temporal area are more likely to change module passing from SC to FC networks.

A deeper analysis of the first five Principal Component is presented in Figure 4. First, we reported in the parameter space, through different colors, the 100 points corresponding to the right tail (≈ 2.5% of the values) of *PC* coefficients, in order to better pinpoint the regions where the *PC* were most expressed (Figure 4(a)). Then, for each component, we only considered the partitions corresponding to these points to investigate which brain areas show the highest variability in the modules’ allegiance between anatomical and functional networks, and across subjects (Figure 4(b,c)). Moreover, for each set of partitions corresponding to the five *PC*, we also reported the agreement matrix and a representative partition for both structural and functional networks (Figure 4(d-h)). The agreement matrix shows the probability that each pair of nodes belongs in the same module in the subspace identified by *PC*_*s*_. *PC*_*s*_ were mostly expressed in different zones of the parameter space and for each of them we could observe different values of community entropy within the ICNs or across subjects, as well as different agreement matrices and representative partitions. For instance, as we intuited before, *PC*_1_ was mostly expressed at low *ω*, and mid-range *γ* and *η*. The partitions falling in this subspace presented a good overlap between structural and functional connectivity. The brain regions whose module’s assignment was most variable belong to the ventral attention network, the DAN and part of the somatomotor area, which, as one can observe in the representative partitions, were joined together in the functional partition, while remaining separated in the structural one. On the contrary, the highest level of inter-subject variability is observed in the brain areas underlying temporal nodes. Similar observations, with different conclusions, can be done for the other four components.

**FIG. 4.**
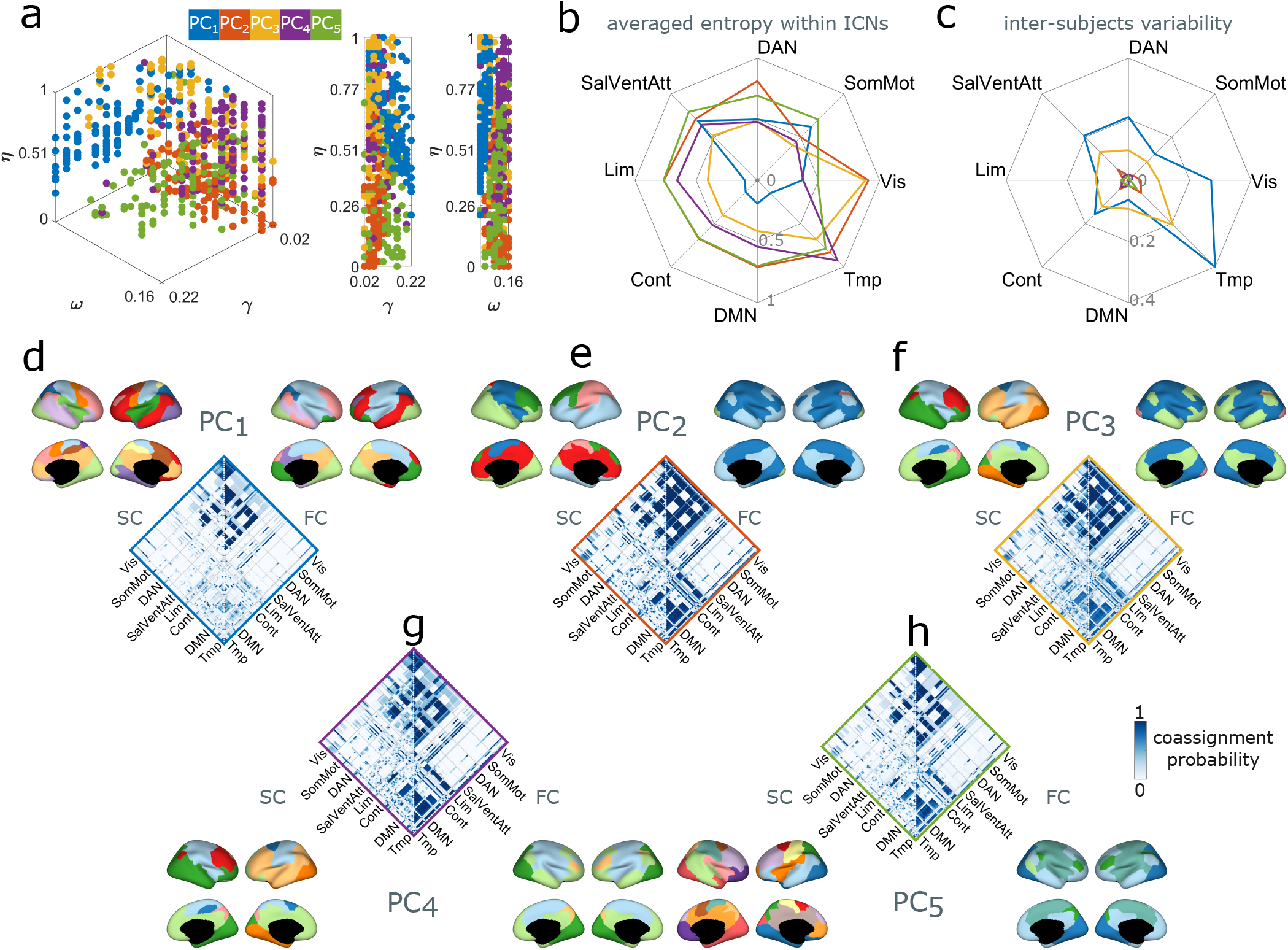
Modes of variability and representative partitions. (a) Projection on the parameter space of the 100 points, corresponding to different combinations of *γ, ω, η*, where the first five components are more expressed. Components are identified through color code. For each component we computed the community entropy between the 100 SC and FC partitions identified in panel (a). By averaging these values across subjects and within functional systems (b) we obtained an information about the nodes whose assignment varies most within each component. Instead, by averaging across combination of *γ, ω, η* of each of the five subspace (c) we gained information about the variability of node’s assignment across subjects, for each component. These results have been reported through spider-plots. (d-h) For each component we selected the partitions corresponding to the points highlighted in panel (a) and we computed a representative partition among them, for both structural and functional networks, and the agreement matrix (co-assignment probability of each pair of nodes).

An analogous PCA analysis has been conducted to evaluate modes of variability of the modular structure across subjects, in both cases of structural and functional connectivity, and it has been reported in the Supplementary Material (Figures S8, S9).

Taken together, these results suggest the existence of inter-modality variability patterns that are well structured in the parameter space. The way sub-systems are coupled between structural and functional connectivity highly depends on the choice of the *γ, ω, η*-values. In other words, the same functional or cognitive system could participate in the same modules in functional and structural networks, or not, according to the topological scale of the partitions (tuned through *γ*) or their consistency across subjects (modulated by *ω*). This information is crucial and has to be borne in mind by a potential user, above all in contexts where structure-function relationships are linked to clinical or behavioral measures of the human brain. Such relationships would be dependent on the choice of *γ, ω* and *η*, which in turn are set arbitrarily by the user.

### Segregation and integration of the modular structure predict inter-modal and inter-subject variability

Previous studies have suggested that the brain’s modular structure helps support segregated (specialized) processing, while the presence of few “hub” nodes helps integrate information across modules. Here, we analyzed the relationship between measures of segregation and integration of the brain networks and inter-modality/inter-subject variability of the modular structure.

As index of integration we used the single-layer modularity score *Q*. Thus, for each subject and point in the parameter space, we computed modularity in anatomical partitions (*Q*_*s*_), functional ones (*Q*_*f*_), and the modularity considering functional modules imposed on the relative anatomical networks (*Q*_*sf*_). We obtained three matrices *NS*(123) × [*N*_*par*_(3723)]. Averaging the structure-function community entropy *H*_*sf*_ across nodes we obtained a matrix of the same dimensions that we could compare with *Q*_*f*_, *Q*_*s*_ and *Q*_*sf*_ through a correlation analysis. The results of this analysis are reported in Figure 5. *Q*_*f*_ revealed to be positively correlated with the entropy between modalities throughout the parameter space (Figure 5(a)), meaning that subjects with functional communities highly modular (well segregated) tend to present greater differences between the structural and functional modular structures. Similarly, *Q*_*sf*_ appeared to be anti-correlated with *H*_*sf*_ (Figure 5(c)) so that, intuitively, the highest is the modular overlap between functional and structural connectivity matrices and the lowest is the cross-modality entropy. These results do not apply for extremely low *γ*, where the correlations were not significant. On the other hand, *Q*_*s*_ was informative of *H*_*sf*_ only for a limited number of combination of {*γ, ω, η*} (see Supplementary Figure S11).

**FIG. 5.**
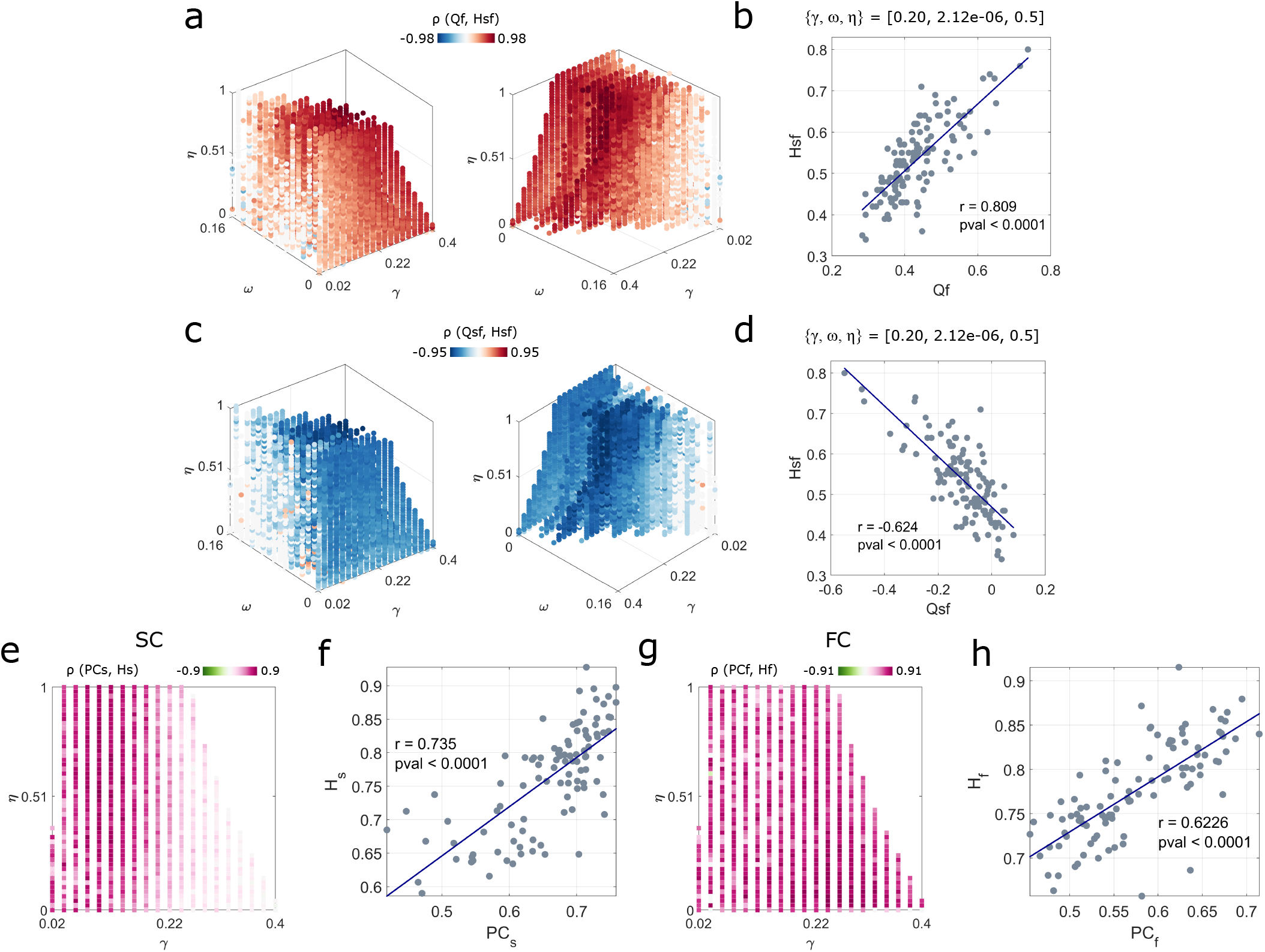
Participation coefficients predict inter-subject variability. (a) Projection on the parameter space of the correlation coefficients of the Spearman correlation computed between the modularity of the functional partitions and the entropy across modality. Non significant values have been set to 0. (b) Example of relationship between *Q*_*f*_ and *H*_*sf*_ at two specific scales. Grey dots represent the values of these variable, while the blue line represents the slope indicated by the correlation. Panels (c) and (d) are an analogue of (a-b) but considering *Q*_*sf*_ instead of *Q*_*f*_. (e) Projection on the *γ, η* space (lowest *ω*) of the correlation coefficients (Spearman correlation) and mean square error computed using the participation coefficient of the anatomical partitions and the entropy of these partitions across subjects. Non significant values have been set to 0. Panel (g) report analogue plots but regarding the participation coefficient of the functional partitions and the entropy of these partitions across subjects. (f) Example of relationship between *pc*_*s*_ and *H*_*s*_ at a specific scales. Grey dots represent the values of these variable, interpolated by the blue line. Panel (h) shows the analogue of panel (f) but for the functional partitions.

To further illustrate that *Q*_*f*_ and *Q*_*sf*_ are strictly linked to *H*_*sf*_ at the subject-level, we focused on a particular combination of *γ, ω, η*. In figure 5(b) we reported the distribution of the single-subject *Q*_*f*_ and *H*_*sf*_ on the *xy*-plane. Those variables were indeed strongly positively correlated (*r* = 0.8, *p <* 0.0001). We observed a similar effect using *Q*_*sf*_ and *H*_*sf*_ figure (5(d)), but this time the two variables were negatively correlated (*r* = − 0.62, *p <* 0.0001).

To measure the integration of nodes in the brain networks we computed the participation coefficient for both structural (*pc*_*s*_) and functional (*pc*_*f*_) networks, obtaining two matrices of dimension [*N* (100) × *NS*(123) × *Npar*(3723)]. We found that this measure is strictly linked to the inter-subject entropy (*H*_*s*_, *H*_*f*_). To show this, we focused on the partitions obtained with *w*_*min*_ = 2.1210^−6^, where *H*_*s*_ and *H*_*f*_ were highest. Once averaged the *pc*_*sf*_ across nodes we computed the Spearman correlation with the *H*_*s*_, *H*_*f*_ matrices. Results are reported in Figure 5(e-h). Both *pc*_*s*_ and *pc*_*f*_ were highly correlated with *H*_*s*_ and *H*_*f*_. In the anatomical networks however, this relationship was attenuated at high gammas.

To see which brain regions participate more in the balance between segregation and integration, we averaged the participation coefficient and the nodal modularity (obtained computing the contribution of each community to *Q*, see Methods) within the ICNs, and we reported their mean values together with their confidence interval through violin plots in Supplementary Figure S12. Among the functional networks, somato-motor, DAN and Salience areas participated mostly in the communication within the modules they belong to (high *Q*_*f*_, low *pc*_*f*_), while DMN, limbic and control network areas showed opposite trends, revealing high integration in the brain network. In the anatomical networks these trends appeared attenuated.

In this section we investigated the relationship between the inter-modality/inter-subjects coupling of the modular structure and two of the best-known measures used to characterize the potential function of nodes given a modular structure. Overall, the results suggested a strong dependence between all these factors. Interestingly, the segregation of functional modules is positively correlated with how much these modules are coupled with the structural ones. We emphasize that these relationships could have been explored only with our extended model. In fact, with the conventional multi-layer framework, we would have lost the subject-dimension on which we built the correlation analysis.

### Detecting multi-modal multi-subject modular structure in the Human Connectome Project dataset

In order to validate the results of the study, we reproduced the analysis on a subset of the Human Connectome Project dataset [103]. It comprises anatomical and functional connectivity data from 95 unrelated healthy adult subjects.

After having built a multi-subject multi-modal network as in Figure 1b, we ran our multi-modal multi-layer modularity optimization algorithm. Then, in order to assess patterns of inter-modality variability, we performed PCA on the entropy matrices obtained computing the node’s entropy between structural and functional partitions in each point of the parameter space given by *γ, ω, η*. We reported in Figure 6(a-e) the projection on the brain surface of the first five principal component’s coefficients, which explained most of the variance of the variability across modality ([12.99, 7.78, 5.73, 4.66]% respectively).

**FIG. 6.**
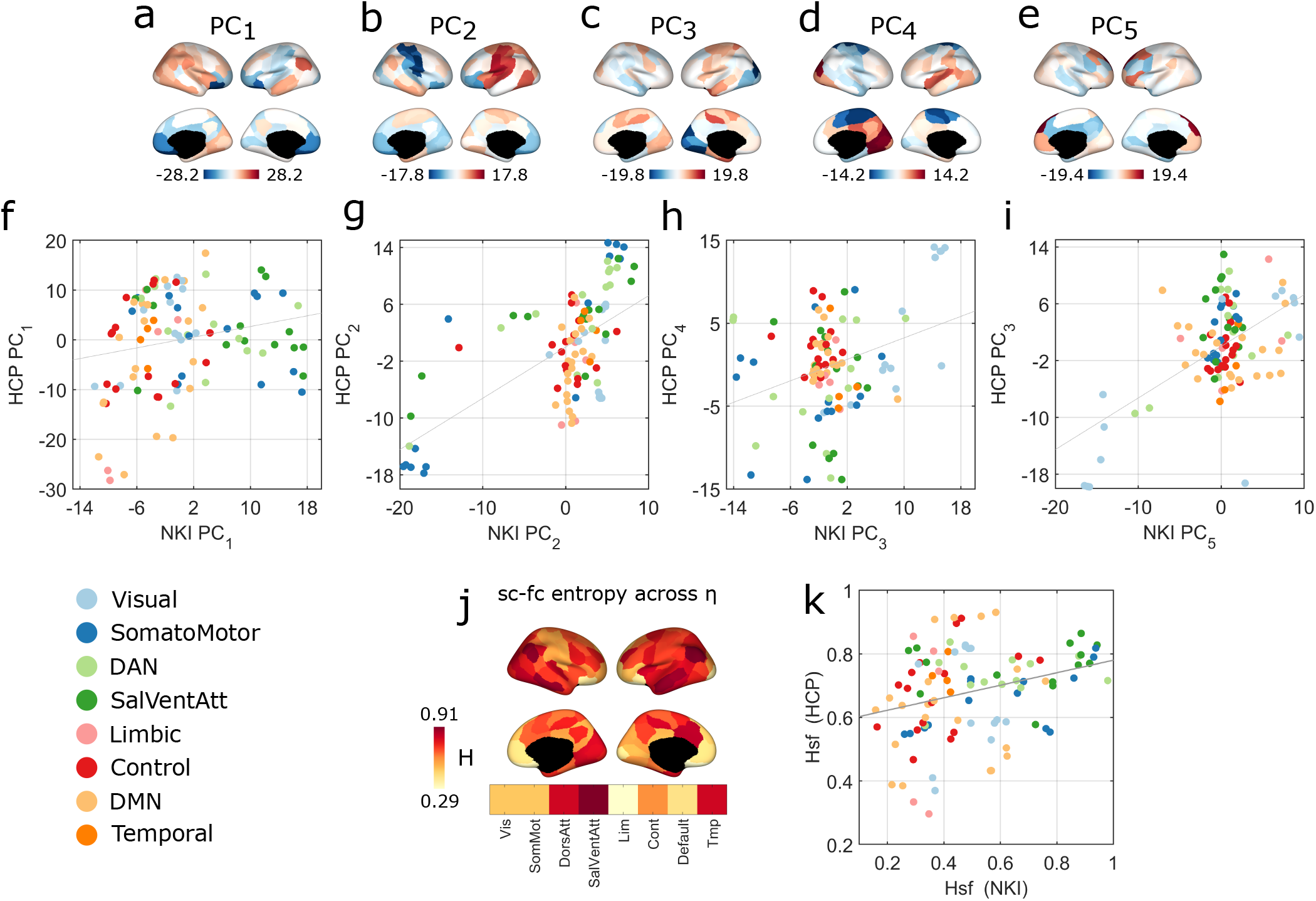
Analysis on the Human Connectome Project dataset. (a-e) Representation on the brain cortex of the coefficients of the first five components generated from the PCA analysis performed on the HCP dataset. These components broadly correspond to those of the NKI dataset, as shown through the correlations maps reported in panels (f-i). (j) Entropy between FC and SC partitions of the HCP averaged across *η* (*γ* = 0.10; *ω* = 0.02) and projected on the brain surface. This is the equivalent to Figure 2(l) for the HCP dataset. At this scale, the obtained values of entropy across modality are correlated across the two different datasets (*r* = 0.3, *pval* = 0.003) (k).

The PCA analysis executed on the NKI and HCP datasets led to comparable results in terms of patterns of entropy between connection modality. To prove it, we looked for statistical correlations between the scores of the first five components of the two datasets. Results are reported in Figure 6(f-i). We found that the first components of the two datasets, 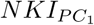 and 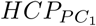, are positively correlated (*pval* = 0.03, *r* = 0.21). For both datasets, in the subspace defined by these components, the brain regions whose modules assignment differs most between structural and functional connectivity belonged to the DAN, the Salience and Ventral Attention networks, and the somatomotor area. At the same time, the DMN together with the Control Network and Temporal areas pertain to modules shared with SC and FC matrices. Note that we have already commented on this for the NKI networks in Figure 3(k). Here we are demonstrating the robustness of the results by validating them on an independent dataset. Other significant correlations, even stronger, have been found between: 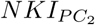 and 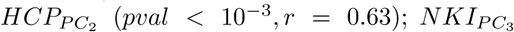 and 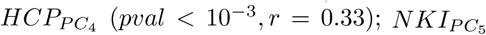 and 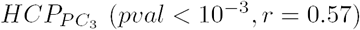.

We further illustrated the consistency of results across independent datasets by focusing on the inter-modality variability at one specific scale (see Figure 6(j,k))). We selected the partitions corresponding to {*γ* = 0.12, *ω* = 0.06 } for the NKI, and {*γ* = 0.10, *ω* = 0.2}. Here, we observed low inter-subject variability and 7 modules for the functional partitions (*CNf*_*NKI*_ = 7 ± 1.9, *CNf*_*HCP*_ = 6 ± 0.9) and 10 for the structural ones (*CNs*_*NKI*_ = 13 ± 1.7, *CNs*_*HCP*_ = 7 ± 0.6). At this scale, the system-averaged values of inter-modality variability significantly correlated in the two datasets (*r* = 0.3, *pval* = 0.003).

We replicated the analysis on a second dataset, the HCP, independently acquired, but similar in terms of participants’ number and age. The reported findings corroborated the results that we first obtained with the NKI dataset. Indeed, modes of inter-modality variability appeared to be robust across datasets and confirmed the multi-scale organization of the human brain networks.

## DISCUSSION

In this work, we explored how the modular structure of the human brain networks is organized across structural and functional connectivity metrics. While most of the literature focuses on the comparison and prediction at the nodes and edge weights level [97], we investigated structure-function relationship at the mesoscale level. In doing this we proposed a novel multi-modal framework, developed on top of the widely-used multi-layer modular optimization [77], where structural and functional connectivity matrices are stacked and linked by three resolution parameters {*γ, ω, η*}. Through this extension we detected the modules of the human brain networks across structural and functional connectivity and among subjects, simultaneously. The three resolution parameters represent a lens through which we could investigate the modular structure of the brain networks, zooming in and out towards organizational traits that are unique or shared among connectivity modalities or subjects. Thus, we explored how modules are configured across spatial scales and at different levels of coupling of the nodes across subjects and types of connectivity.

### Relationships between structural and functional modular structure across subjects

The principal aim of this work was to characterize the community organization of brain networks across types of connectivity. Our results suggest that there is not a single way to describe the overlap between structural and functional modular organization. Rather, we observed that this overlap depends on the spatial resolution of modules (controlled by the parameter *γ*), but also on the inter-subject and inter-modality resolution (regulated by the parameters *ω* and *η*, respectively). Trivially, the strength of the inter-modality resolution (*η*) was proportional to the overlap between structural and functional modules. As for the dependence on the spatial resolution (*γ*), previous studies have shown how brain networks display a multi-scale community organization [11]. Therefore it is intuitive that the relationship of this organization between different types of connectivity will also be multiscale. Specifically, we observed a higher SC-FC modular coupling in finer partitions. Tuning the cross-subject resolution (*ω*) instead, we found a higher coupling between structural and functional partitions when considering low community variability across subjects.

In this study, we quantitatively found, through principal component analysis, modes of variability between anatomical and functional modules. We described in detail the structure-function relationship explained by the first five components, expressing most of the variance in the *PCA*, and located in the parameter space as follows: *PC*_1_ high-*η*, low-*ω*, high-*γ*; *PC*_2_ low-*η*, low-to-high-*ω*, low-*γ*; *PC*_3_ low-*γ*, low-*ω*, high-*η*; *PC*_4_ high-*η*, high-*ω*, low-*γ*; *PC*_5_ high-*γ*, high-*ω*, low-*η*. These results confirmed that the structure-function coupling, at the modular level, is mediated by the topological scale. In the best possible condition from the SC-FC coupling point of view, represented by *PC*_1_, the regions whose community assignment varies the most between modality are mainly associated to the ventral attention network. Interestingly, this might align with results in [105], where the coupling between structural and functional connectivity in this area has shown to be statistically lower than chance when compared to a null model. The DMN instead, is the functional system where the relationship between structure and function is more heterogeneous, as for some components we observed a coupled modular organization, while for some other components we did not. This could be due to the high number of circuits in which DMN participates and the variable dynamics it exhibits [21, 104, 108]. In *PC*_2_, high entropy values between structural and functional communities are widespread in the cortex, involving all the ICNs except for the somatosensory system. This result fits recent studies where structural and functional connectivity was found to be more consistent in the unimodal (sensory) cortex with respect to transmodal cortex [81, 84]. A possible interpretation for the general high-entropy in *PC*_2_ could be linked to the fact that it is expressed at low-*ω*, where partitions are variable across subjects. As functional brain networks have been shown to be subject-specific [34, 46, 62], so will be their relationship with the underlying structural network. Thus, when the coupling among subjects is low (low-*ω*), it is reasonable to expect a high variability between structural and functional partitions over different brain regions.

All these findings suggest that the relationship between anatomical and functional modular organization is not well summarized by a single spatial or temporal scale. Different studies have been carried out focusing on a single-scale community structure and its relationship with cognitive output, providing important insight into brain functions. However, our work, together with [11, 17], suggests that multi-scale analysis are needed to gain a more comprehensive understanding of the relation between brain structure, function and cognition.

### Methodological innovation

In this work we extended the classical multi-layer modularity maximization framework to address a long-standing question in neuroscience: what is the relationship between anatomical and functional modular organization and how this relationship is expressed across different individuals? Exploiting the flexibility of modularity optimization [28], we built a modularity matrix that contains multi-modal and multi-subject networks contemporaneously. The advantage of this method consists of having communities matched across layers, representing either individuals and type of connectivity. In this way, it allows a straightforward analysis of the modular structure: to get an estimate of whether communities are the same across connection modality and/or subjects, one can trivially compare community labels, without the need for additional heuristic. Previous studies already used the multi-layer framework [25, 102] to track communities across subjects [17], time [9, 15, 18, 92], learning paradigms [7, 43], tasks [22] clinical cohorts [53], or to model brain dynamics [60]. Here, for the first time, thanks to the proposed extension, we could observe how the modular structure reorganizes across two distinct directions, that in our case were subjects and connection modality.

Ultimately, where is the novelty of our approach? Why it is useful? And why would we want a multi-modal, multi-subject model? The main novelty lies in that it allows a straightforward comparison of the modular structure across multiple domains (in our case subjects and connectivity modality). Not only this is useful for discovery purposes (i.e. the investigation of structure-function relationship in the healthy brain), but also it could be applied in context with high clinical relevance, e.g. in clinical populations, where features of the relationships between structure and function can be used as biomarkers. An example could be a structure-function investigation in stroke patients, where anatomical connectivity is damaged and the way its relationship with functional connectivity mutates after the stroke event is crucial for rehabilitation purposes. Furthermore, one could also consider a multi-modal analysis across different tasks or cognitive states, where we might expect a variation in the structure-function coupling. Thus, this framework opens the door to new studies in which multiple co-occurrent factors can be taken into account in the analysis of the human brain topological organization. In general, more multi-layer approaches are needed to incorporate multiple channels of connectivity, to obtain a more thorough knowledge on the brain functioning.

Another important methodological aspect regards how we treated structural and functional matrices before incorporating them into the modularity matrix. In fact, it is good practice building multi-layer networks where layers are made of comparable weights, otherwise layers with higher weights may impact excessively the modularity optimization and thus the communities estimates. It is worth noticing that structural and functional networks are two distinct mathematical objects. A structural network is a sparse matrix with only positive weights, while a functional network is a full correlation matrix with both positive and negative weights. To overcome this difference, we converted structural matrices into structural *correlation* matrices. In this way, both functional and structural networks could be entered in the modularity matrix as similarity matrices, with weights falling in the same range, without running the risk of having one dragging the other in the community detection. This conversion also allowed us to use the same null model in the modularity optimization, for both structural and functional layers. Finally, we want to point out that our computing the pairwise correlation between rows of structural matrices (to obtain correlation matrices) is similar to the computation of the matching index [55], which captures the overlap of connection patterns for each pair of vertices. This index has been widely employed in the recent network neuroscience literature, for instance to study connection fingerprint [95], build generative models [14], or predict functional connectivity patterns from measures of network communication [48].

### Limitations

In this study we focused on a specific kind on communities, i.e. modules, that are defined upon an assortative criterium [44]. It is well established that they well represent the brain networks organization [94], promoting states of segregation and integration [40], necessary for an efficient brain functioning [106]. However, brain networks can also present different kind of organizations, based for example on core-periphery structure, disassortative communities or diffusion models [16, 31, 32, 78]. Recent studies started to explore different levels of brain networks organization [33], with overlapping modules, so that future advancements could extend these efforts toward a multi-layer modeling. Furthermore, modularity maximization is not the only way, nor the best one, to find assortative communities in networks. However, as discussed in [28], it is a useful tool to address specific questions of neuroscience, like the structure-function relationship, in a principled manner.

All the comparisons between structural and functional partitions, or among subjects, have been made through the Variation of Information. We have chosen this index in line with previously mentioned studies based on modularity optimization. However, there are a number of indices that can be used alternatively [42].

Finally, we tested our model on a single parcellation of the brain networks. We used a 100 nodes parcellation to save computational resources. In general, multi-layer approaches are computationally expensive. In fact, while in single layer networks operations like modularity maximization are made on a modularity matrix whose dimension is given by the number of nodes, in multi-layer networks this same dimension is multiplied by the number of layers. Our proposed approach, by doubling the dimension of the multi-layer modularity matrix, requires even more memory and computational power and thus, does not scale well for extremely large datasets. Using a 100 nodes parcellation we optimized an already big modularity matrix, of dimension 24,600 ([123 subjects × 2 modalities × 100 nodes]). However, further analysis can be done to validate our results on finer parcellations of the human brain cortex [3].

### Future advances

There exists an increasing body of literature bringing attention to the influence of the brain network architecture on cognition and behavior [20, 67, 74], and previous studies have shown that structure-function relationship and human behavior are also related [69]. Moreover, structure-function relationship characterizes subjects individuality [50]. In this work, we propose as a supplemental investigation a preliminary analysis of how measures derived from four different cognitive assessments can be associated to the five modes of variability between structural and functional modular organization (paragraph S8 in Supplementary Material). We observed that a relationship exists, and it is scale-dependent. Indeed, across the five principal components, most of the brain regions showed different patterns of correlation between SC-FC community entropy and IQ-based measures. Notably, the DMN is the subsystem whose patterns of correlations vary the least, maintaining always a positive correlation between IQ and its entropy across structure and function.

However, we only presented a proof-of-concept analyses to relate multi-scale partition information to cognitive assessments. To conclusively establish such relationships would likely require much larger sample sizes [65] and the application of additional tests and methodologies [107]. For these reasons, we presented this part of the work without drawing strong conclusions (and we also invite the reader to make the same). Rather, we illustrated brain-wide patterns at different scales, proving that our proposed methodology, by preserving single-subject information while allowing the exploration along a third dimension, could be useful in future studies to properly investigate brain-behavior associations.

Finally, we want to highlight that our focus were on a restricted category of participants, that is healthy adults. Future works could exploit the proposed methods and extend the study to lifespan studies or clinical cohorts, to investigate if and how the relationship between structure and function evolves across ages or diseases.

## CONCLUSION

In conclusion, we developed a novel multi-layer framework able to incorporate multiple modes of connectivity, so that we could quantify how much the network modular structure varies across connection modality and subjects simultaneously. We investigated the relationship between structural and functional brain networks modular organization, and how it is shaped across subjects and at different spatial scales. While confirming previous findings on specific brain areas, our results provide also evidence that these relationships are recapitulated by scale-dependent modes of variability. Overall, this work not only increases the state-of-the-art methods and knowledge about structure-function relationship, but also enables new analysis that can be done to comprehensively map the human brain networks organization across multiple domains.

## CODE AVAILABILITY

MATLAB code to build the multi-modal and multi-subject network and run the extended modularity optimization is available at https://github.com/mariagraziaP/multimodal_multisubject_modularity.

