## Supplementary Material for "Multi-modal and multi-subject modular organization of human brain networks"

### Supplementary Information: Multi-modal multi-subject modular organization of the human brain network

#### Contents

**S1. Schematic representation of the methods** **2**

**S2. Preliminary statistics on communities** **4**

**S3. Variability between structural and functional modular organization at specific scales** **7**

**S4. Explained variance of the principal components** **9**

**S5. Principal Component Analysis – patterns of inter-subject variations** **10**

**S6. Variability of community structure across subjects at specific scales** **13**

**S7. Segregation and integration of the network modules** **15**

**S8. Brain-behavior correlation** **17**

**S9. Brain-behavior correlation considering the inter-subjects community entropy** **21**

### S1. Schematic representation of the methods

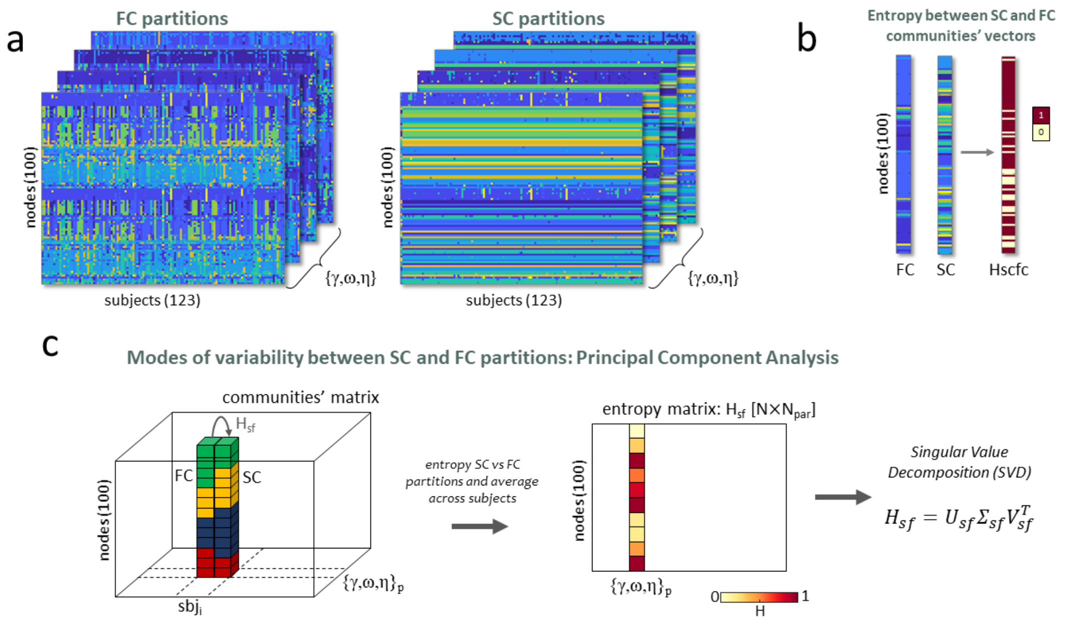


**Figure S1: Schematic representation of the steps to perform the Principal Component Analysis.** (a) Output of the multisubject multimodal community detection framework. We obtain a partition of both structural and functional networks into modules for each subject and each combination of resolution parameters $\{\gamma, \omega, \eta\}$. (b) Example of community entropy computed between two partitions. It returns 1 if the node belongs to the same cluster partition-wise, and 0 otherwise. (c) We build an entropy matrix $H_{sf}$ by computing the community entropy between SC and FC partitions for each subject and each $\{\gamma, \omega, \eta\}$, and then averaging across subjects. We decomposed the resulting entropy matrix in components through the Singular Value Decomposition.

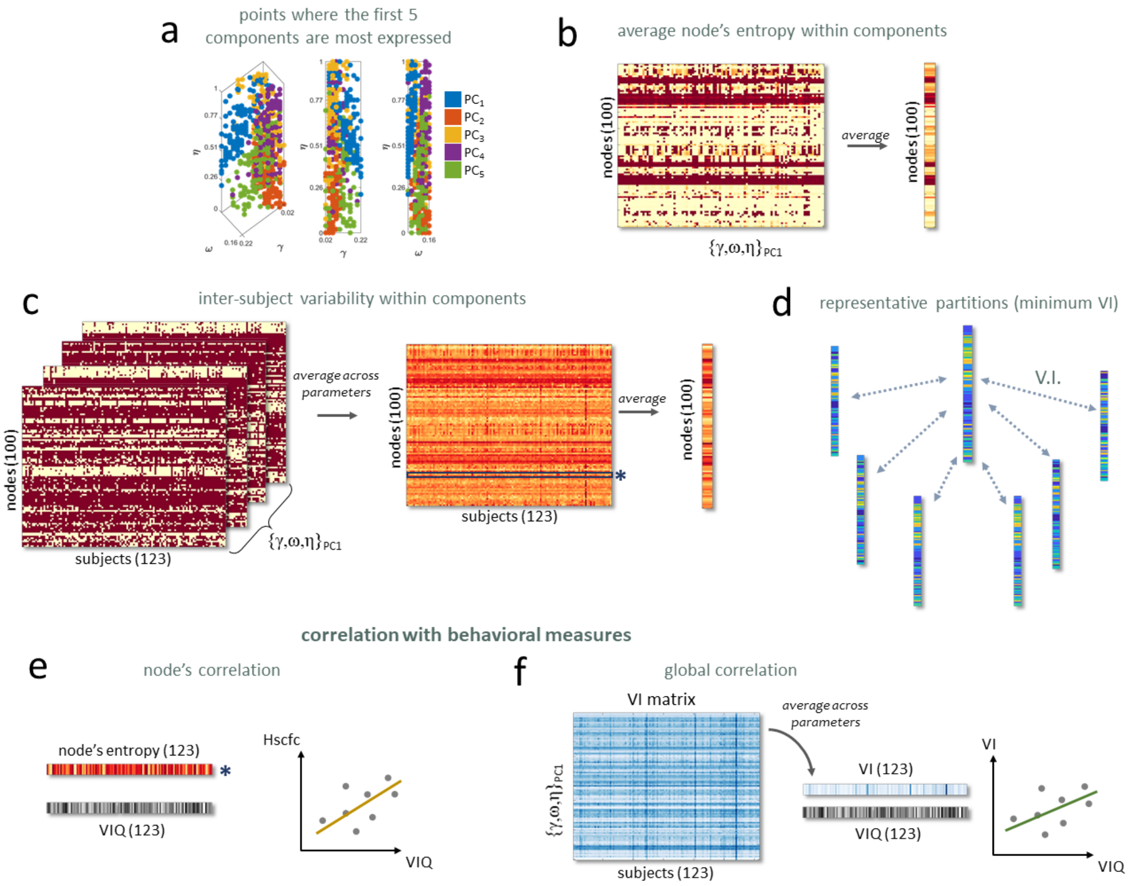


**Figure S2: Analysis of the principal components.** (a) Representation, on the parameter space identified by $\{\gamma, \omega, \eta\}$, of the highest 100 coefficients, for each one of the first five components. (b) For each component, we computed the community entropy between SC and FC partitions corresponding to the 100 combinations of $\{\gamma, \omega, \eta\}$ identified in panel a, and we averaged across subjects, obtaining an entropy matrix of dimension [nodes x 100]. Then by averaging the rows of this matrix we obtained a measure of the node's entropy, and the mean entropy in the ICNs, within the component, that we represented in Figure 3b. (c) For each component, we computed the community entropy between SC and FC partitions corresponding to the combinations of $\{\gamma, \omega, \eta\}$ identified in panel a, and this time we average across parameters, obtaining a matrix of dimension [nodes x subjects]. Then, by averaging the rows of this matrix we obtained a measure of the entropy of the nodes (or ICNs) among subjects, that we reported in Figure 3c. (d) Among a set of partitions we consider the centroid as the representative one. We compute the Variation of Information (VI) between each pair and select the partition with lowest VI. (e) For each component, from the entropy matrix computed in panel c we extracted the rows, indicating a subjected-level measure of the community entropy between SC and FC partitions. We computed the Spearman correlation between each rows and IQ indices, obtaining for each component a node-level information about the relationship between the SC-FC community entropy and behavioral measures. (f) We also obtained a global measure of the correlation between SC-FC community structure and behavioral indices. For each component we computed the VI between SC-FC partition taken among the 100 combinations of $\{\gamma, \omega, \eta\}$ identified in panel a, obtaining a VI matrix of dimension [100 x subjects]. By averaging column-wise we obtain a row containing a subject-level measure of the VI, that we spearman-corelated with the IQ indices.

### S2. Preliminary statistics on communities

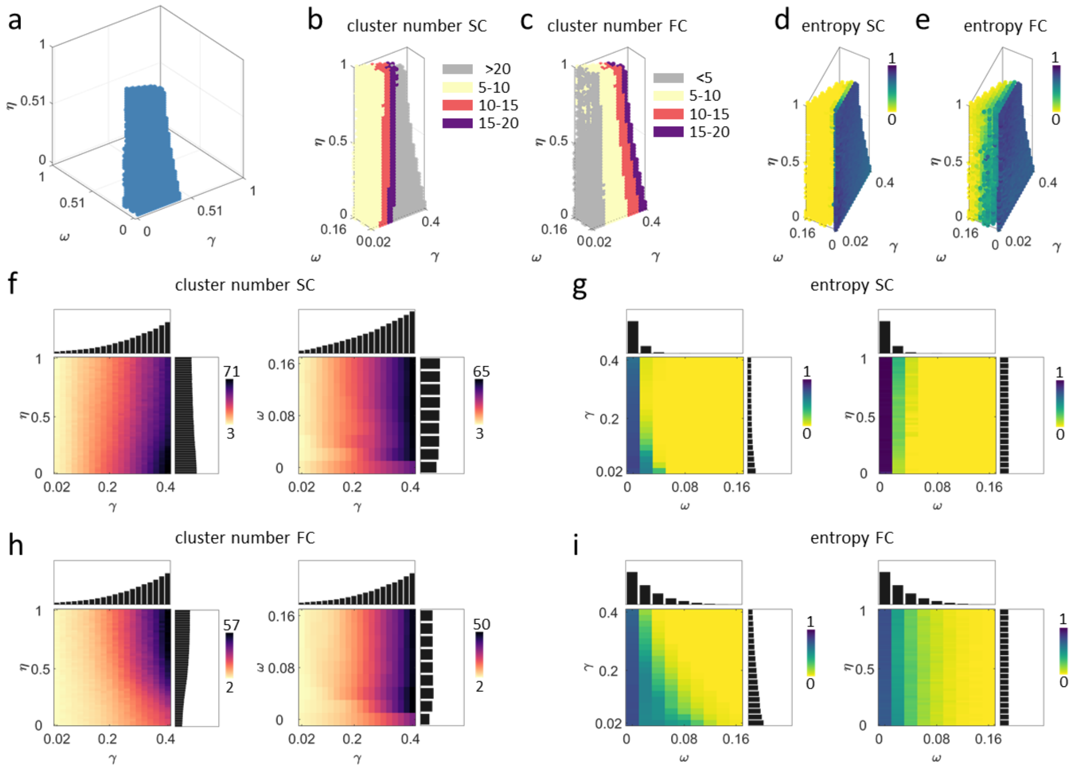


**Figure S3: Statistics in the parameter space.** (a) Representation on the parameter space of the combinations of $\{\gamma, \omega, \eta\}$ leading to physiologically meaningful partitions. These points have been selected with criteria based on number of clusters and community entropy (see Methods). (b,c) In each non-trivial partition we computed the number of cluster and reported it through color-code for both structural and functional networks. The clusters number is mainly affected by the $\gamma$-value so that lower $\gamma$s lead to partitions with fewer clusters and vice-versa (f,h). Functional networks tend to be parsed in fewer clusters with respect to the structural ones. (d,e) In each non-trivial partition we also computed the community entropy among subjects. This measure is mainly affected by the $\omega$-value so that higher $\omega$s lead to partitions more consistent across subjects (g,i). Functional partitions tend to be more variable across subjects with respect to the structural ones.

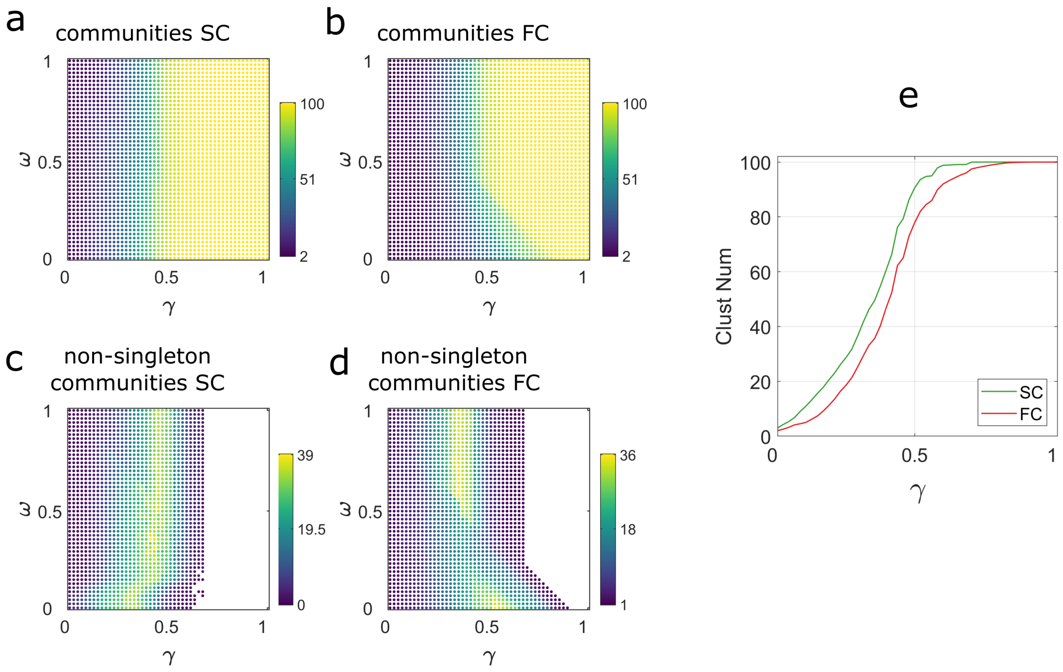


**Figure S4: Statistics in the parameter space for the preliminary group-average analysis.** In each structural (a) and functional (b) partitions we computed the number of clusters, and reported this measure through a color-code on the parameter space defined by $\gamma$, $\omega$. Clusters number is mainly affected by $\gamma$ in both types of connectivity, so that higher $\gamma$ lead to finer community structure. We also computed the number of non-singleton communities, i.e. communities composed by only one node, and reported this measure in panels (c,d). Here, blank spaces correspond to combinations of $\gamma$, $\omega$ that render partitions with number of lcusters equal to the number of nodes (i.e. all singleton communities). In panel e we show the trend of the number of clusters with respect to $\gamma$, averaging along the $\omega$ dimension.

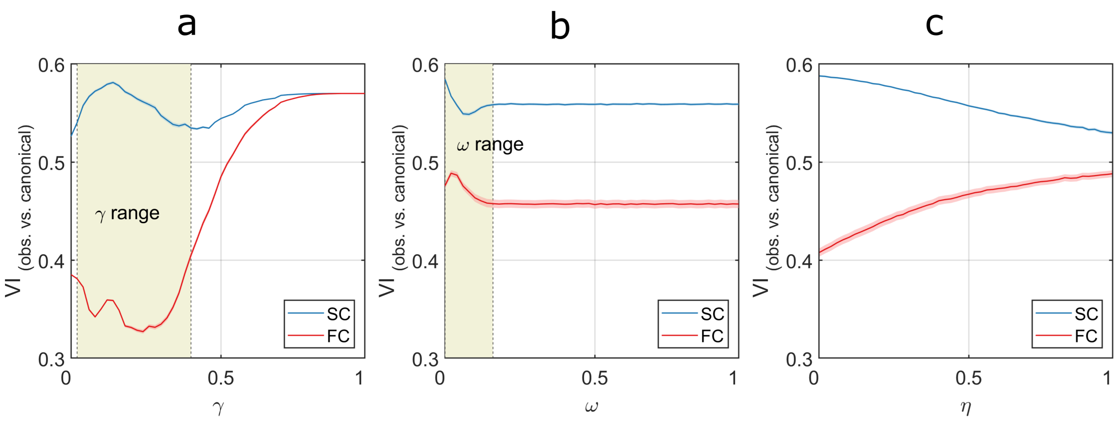


**Figure S5: Distance between the discovered communities and the canonical systems.** We represent the distance - in terms of VI - between both functional and structural communities discovered all across the parameter space and the YEO canonical systems. We show the trend of VI with respect to $\gamma$ (panel a), $\omega$ (panel b), and $\eta$ (panel c). In each panel, we overlap in the background the range of the resolution parameters chosen according to our criteria based on number of clusters and inter-subject entropy (see Methods section). First of all, we noticed that functional partitions tend to be more similar the canonical systems with respect to structural ones, since they present lower VI values. This is trivially explained by the fact that the canonical systems have been defined from functional networks. Second, we found that this similarity is mostly influenced by $\gamma$ (in panel a VI spans a wider range). Notably, the chosen interval for this $\gamma$ corresponds to the space where discovered and functional partitions are more similar. Finally, we observed that in the selected $\omega$-range VI values are higher than in the on-selected ones (panel b). However, this bump is likely caused by the inter-subject variability that we wanted to preserve in our analysis. Moreover, this initial "bump" of VI is extremely contained with respect to the whole range occupied by VI across the parameter space. All these considerations on panels a and b validate our selection criteria. As for panel c, where we show the trends of Vi with respect to $\eta$, it further validates our framework. In fact, it demonstrates how increasing $\eta$ leads to a progressive coupling between FC and SC partitions. The former is maximally similar to the canonical systems for low $\eta$ (that is in the standard condition in which they have been introduced). Then VI increases, meaning that functional partitions progressively diverge from those systems to create a better fit to the structural ones. The contrary happens for SC partitions. They are maximally distant from canonical systems for low $\eta$, but as $\eta$ increases they get closer, in order to better fit the functional ones.

### S3. Variability between structural and functional modular organization at specific scales

Through the novel multilayer modularity formulation, we were able to identify partitions of the brain networks at different spatial scales, and differently coupled across subjects and type of connectivity. However, this method also allows to focus on a restricted combination of $\{\gamma, \omega, \eta\}$ deemed interesting, in order to provide a deeper characterization of communities. Here, we explored how SC and FC partitions reconfigure across $\eta$, getting more and more coupled, focusing on the subset of parameters $\gamma = [0.06, 0.12, 0.20]$, $\omega = 0.06$. This $\omega$-value ensured consistency across subjects, while the three $\gamma$-values enabled to look at the modules at different spatial scales, from a coarser to a finer spectrum of communities.

Results are reported in Figure S6. The three rows of the figure (panels c-n) correspond to the three $\gamma$-values. For these combinations of $\{\gamma, \omega\}$ we observed how the community entropy between SC and FC partitions (averaged for all the subjects) varies across $\eta$ (panels c, g, k). In order to identify the brain areas with highest (lowest) community entropy values, we also represented the mean entropy value across $\eta$ on the cortex surface and within the ICNs (panels d, h, l). Then, we show the results of the correlation between the community entropy and $\eta$ for each node, reporting significant values ($p < 0.05$) of the correlation coefficients (panels e, i, m). Finally, we report the representative partitions at three progressive values of $\eta = [2.1210^(-6), 0.48, 0.99]$, for both structural and functional networks (panels f, j, n).

According to these results, different brain regions exhibit different patterns of community entropy across $\eta$ and $\gamma$. Increasing values of $\eta$ results in low entropy between FC and SC partitions. At the same time, also setting the $\gamma$ to obtain finer partitions resulted in high coupling between FC and SC partitions even at medium $\eta$-values. On the contrary, at coarser scales (corresponding to $\gamma = [0.04; 0.12]$) we can observe how some regions keep different module's assignment even with high $\eta$-values. For $\gamma = 0.06$, the nodes involved in the temporal and visual areas, and those in the ventral and dorsal attention networks, show the lowest coupling between FC and SC partitions all over the $\eta$-range. At the scale defined by $\gamma = 0.12$ we can observe an increasing tendency of some brain areas to overlap more and more between FC and SC as $\eta$ increases. These areas involve the limbic system, the control network, the DMN and the temporal area. The ventral and dorsal attention networks are still the regions showing higher community entropy. When $\gamma = 0.2$ all the brain regions showed high modules consistence between FC and SC networks for high $\eta$-values. Globally, in every considered spatial scale, the nodes belonging to the salience and ventral attention network were those whose cluster's assignment remain more variable across modality.

Looking at the representative partitions obtained with these combinations of $\{\gamma, \omega, \eta\}$, we have a confirmation of what just inferred. Increasing $\eta$-values result in highly coupled SC and FC partitions. This coupling turned into a perfect match for high $\gamma$-values. For lower $\gamma$ however, the nodes involved above all in the medial area and the salience network belong to different clusters in the two type of connectivity.

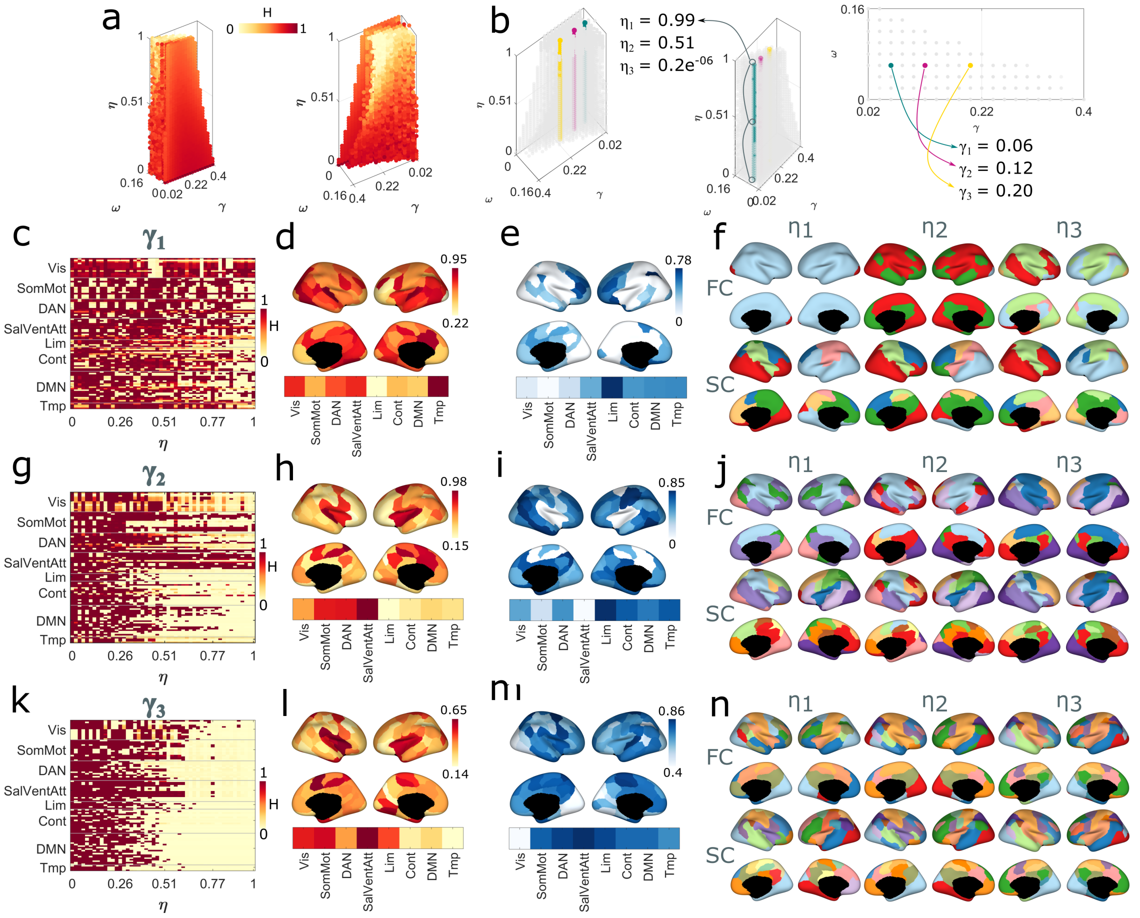


**Figure S6: Modes of variability and representative partitions.** (a) Projection on the parameter space of the 100 points, corresponding to different combinations of $\gamma, \omega, \eta$, where the first five components are more expressed. Components are identified through color code. For each component we computed the community entropy between the 100 SC and FC partitions identified in panel (a). By averaging these values across subjects and within functional systems (b) we obtained an information about the nodes whose assignment varies most within each component. Instead, by averaging across combination of $\gamma, \omega, \eta$ of each of the five subspace (c) we gained information about the variability of node's assignment across subjects, for each component. These results have been reported through spider-plots. (d-h) For each component we selected the partitions corresponding to the points highlighted in panel (a) and we computed a representative partition among them, for both structural and functional networks, and the agreement matrix (co-assignment probability of each pair of nodes).

### S4. Explained variance of the principal components

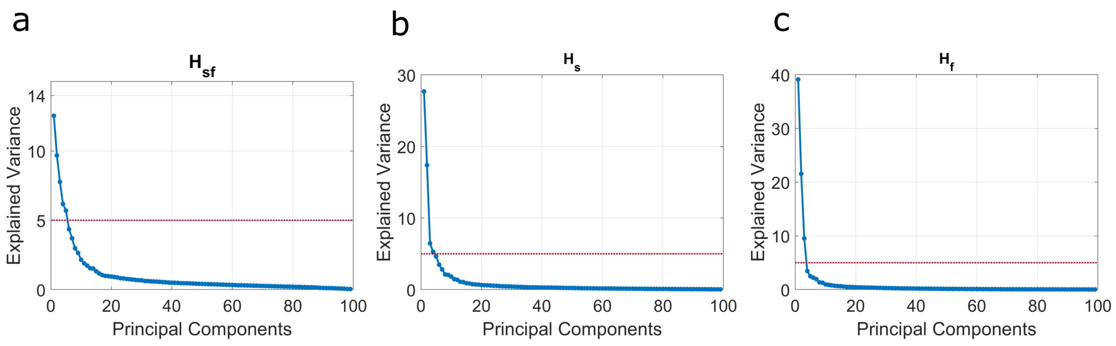


**Figure S7:** Explained variance of the principal components computed on the inter-modality modules' entropy $H_{sf}$ (a), inter-subject structural modules' entropy $H_s$ (b), and inter-subject functional modules' entropy $H_f$ (c))

### S5. Principal Component Analysis – patterns of inter-subject variations

To investigate the modes of variability across subjects (within modality) we built the $H_{sbj}$ matrix (100×3732) concatenating the 3732 vectors (one for each point in the parameter space) containing the normalized entropy values between of the FC (or SC) partitions across the 123 subjects. We report here results relative to the first three components, that, out of the 99 explained most of the variance (components from 4 to 99 explains less than the 5% of the variance, see Fig. S7(b,c), for both analyses based on structural and functional connectivity).

Results are shown in Figure S8-S9. In both case of SC and FC, the coefficients of $PC_1$ observed in the parameter space are most expressed at $\omega$-values where partitions across subjects are fairly consistent. For both FC and SC, the correspondent $PC_1$ scores show that the regions with more variable community assignment across subjects belong to the visual and temporal areas, while the somatomotor area is the most consistent. With respect to FC pattern, the SC one shows that also the limbic system and the DMN are responsible for high entropy. Other patterns of variability can be observed across the other PC. In $PC_2$ and $PC_3$, the coefficients are more expressed in the parameter space where the $\omega$ is the lowest, meaning where partitions across subjects are variable. As expected, the $PC_2$ and $PC_3$ scores indicate regions globally more variable with respect to $PC_1$ scores (Figure S5a,b). The difference between $PC_2$ and $PC_3$ is that are located at the extremes of the $\gamma$ range. Results suggest that for $PC_2$, at low $\gamma$, in both SC and FC networks, DMN and control networks are subject-specific, while the visual and the somatomotor area are more consistent across individuals. SC and FC differ for the community entropy linked to the temporal area. Regarding $PC_3$, SC and FC present more different trends.

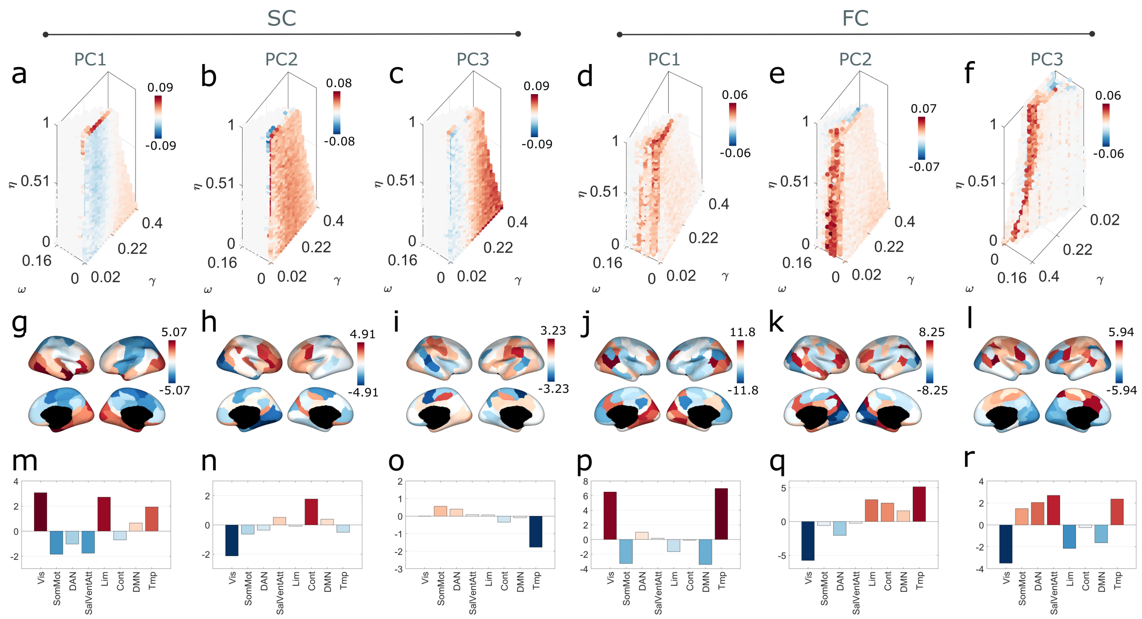


**Figure S8: Principal Component Analysis for the detection of modes of variability across subjects, for both structural and functional connectivity networks.** In the first three columns are reported the results of the PCA for the first three components obtained analyzing structural networks, while in the last three columns those related to functional networks. (a-f): projection of the first three PC coefficients into the parameter space identified by the parameters $\{\gamma, \omega, \eta\}$. (g-l): projection on the cortex surface of the first three PC scores. (m-r): average within the functional systems of the first five PC scores. In each component, brain areas colored with red present community assignment highly variable across subjects. These variations mostly occur in the space of parameter identified by red dots. Blue-colored brain areas instead, identify stable node's community assignment in the relative blue points of the parameter space.

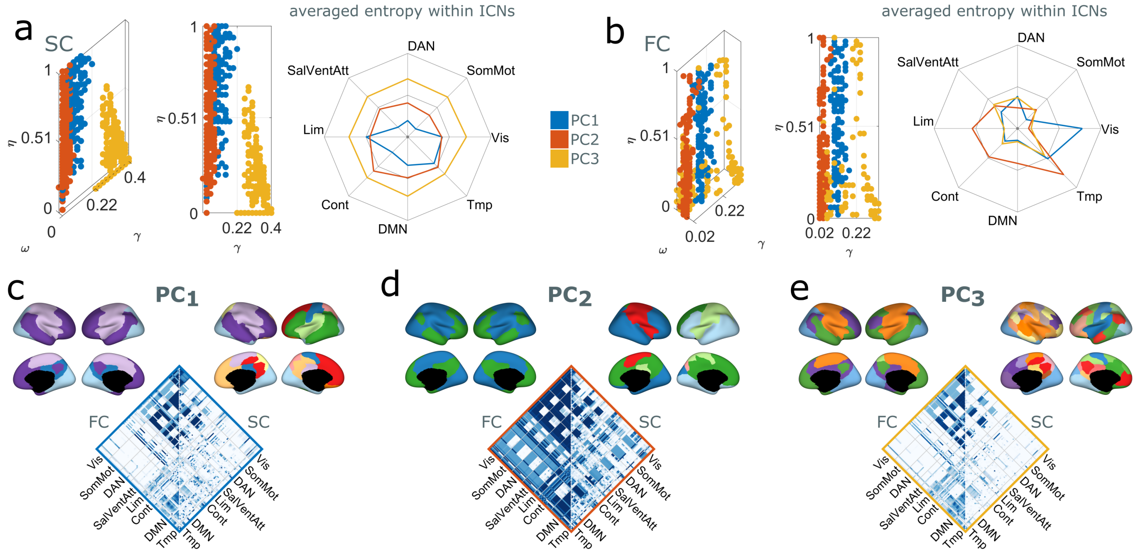


**Figure S9: Modes of variability and representative partitions.** (a) Projection on the parameter space of the 100 points, corresponding to different combinations of $\{\gamma, \omega, \eta\}$, where the first three components of the PCA related to structural connectivity networks are more expressed. Components are identified through color code. For each component we computed the community entropy between the 100 SC partitions. By averaging these values across subjects and within functional systems we obtained an information about the nodes whose assignment varies most within each component. We represent this result though a spider plot. (b) Same results of panel a but related to functional connectivity networks. (c-e) For each component we selected the partitions corresponding to the points highlighted in panels a and b and we computed a representative partition among them, for both structural and functional networks, and the agreement matrix (co-assignment probability of each pair of nodes).

### S6. Variability of community structure across subjects at specific scales

We evaluated how the partitions vary across subjects focusing on a restricted number of $\{\gamma, \omega, \eta\}$. We choose $\gamma = \{0.06, 0.12, 0.2\}$, $\omega = 0.02$ and $\eta = 0.48$ (Figure S10). At this $\omega$ and $\eta$ scales SC and FC partitions are at an intermediate level of coupling, and the partitions across subjects are not totally disjoint nor totally matched, as would it be by choosing lower and higher $\omega$-values respectively. The values of $\gamma$ have been chosen with the same criteria explained in the main paper. For these combinations of $\{\gamma, \omega, \eta\}$ we reported for each subject the mean value of the entropy computed between its partition and the partitions of all the other subjects (panels c and d). In order to identify the brain areas with more (less) variable community assignment across subjects, we reported the entropy value on the cortex, and within the ICNs. Finally, we reported a representative partition for each one of the considered combinations of $\{\gamma, \omega, \eta\}$ for both FC and SC networks (panels d and e).

Results suggest that the coupling of the partitions across subjects is dependent not only by the chosen temporal scale ($\omega$) but also by the spatial scale ($\gamma$) at which we are looking communities. It is also evident that SC networks are globally more stable at the subject-level with respect to FC networks, which are more subject-specific. When $\gamma = 0.06$ the temporal and visual areas, and the nodes pertaining to the control network are highly variable across subjects in the FC networks, while the DMN remains quite consistent. The anatomical networks of the same subjects instead, show high variability in the clusters' assignment of the nodes of the control network, DMN, salience network and the temporal areas. Increasing $\gamma$ to 0.12 we observe different pattern of variability. In FC networks the temporal, visual and limbic areas and the DAN and control networks constitute the clusters less consistent across subjects, while in the SC networks these clusters have been localized mostly in temporal, limbic and visual areas and in the DMN. At a finer scale ($\gamma = 0.2$) the SC networks show partitions with strong correspondence among subjects, except for the visual area. The FC networks instead, still show a variability of the modules, primarily localized in the nodes of the temporal areas, and in those of the DAN and ventral attention networks. Over all the scales, the part of the cortex that is more subject-specific at the modular structure level, for both kind of connectivity is the temporal area.

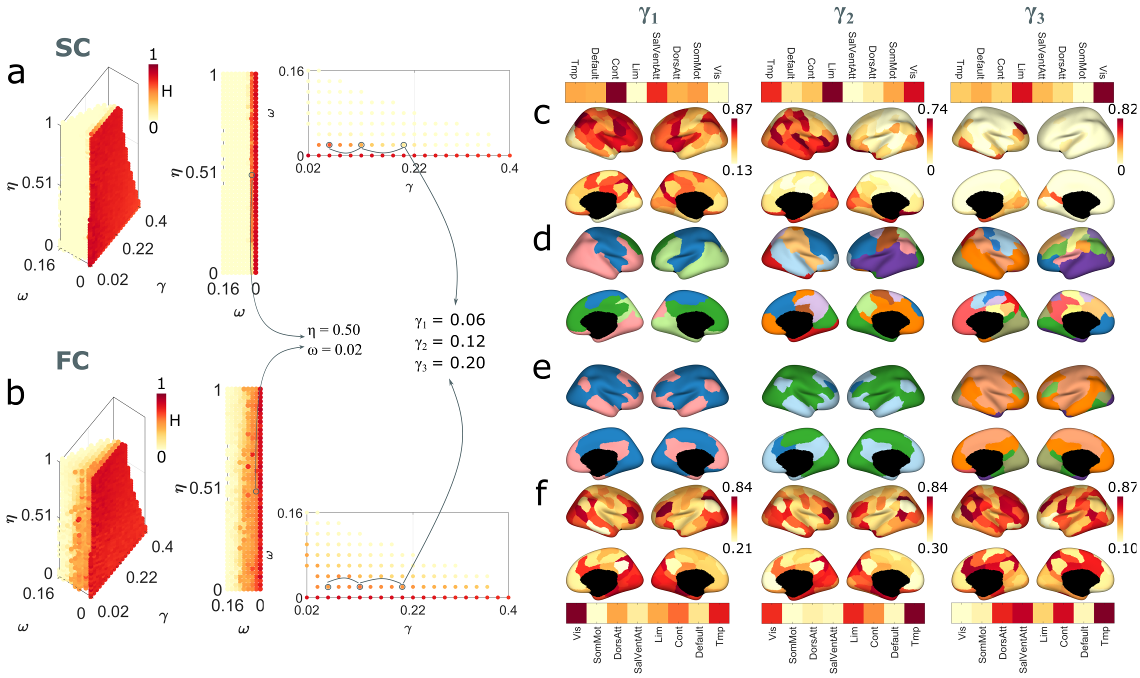


**Figure S10: Reconfiguration of the modular structure across subjects.** (a-b) Projection on the parameter space of the community entropy computed across subjects in the SC and FC networks respectively. We show three views of the parameters space. Yellow points correspond to null entropy, that is coupled partitions, while the contrary is true going towards red colors. (c) Projection on the cortex of the community entropy computed across subjects within the SC partitions for three increasing $\gamma$-values, at $\omega$=0.02 and $\eta$=0.5. (d) Representative SC partitions, among the subjects, for the three values of $\gamma$. Panels (e) and (f) are equivalent to panels (c-d) but related to the functional networks.

### S7. Segregation and integration of the network modules

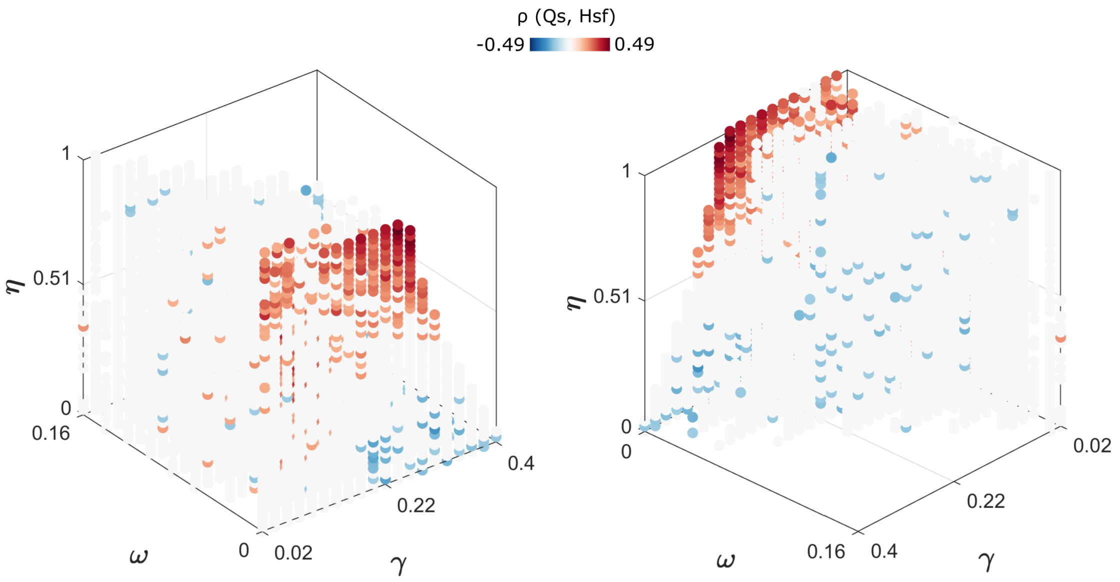


**Figure S11: Relationship between modularity of structural networks and cross-modality entropy.** Projection on the parameter space of the correlation coefficient resulting from the Spearman Correlation computed between the modularity of the anatomical partitions ($Q_s$) and $H_{sf}$

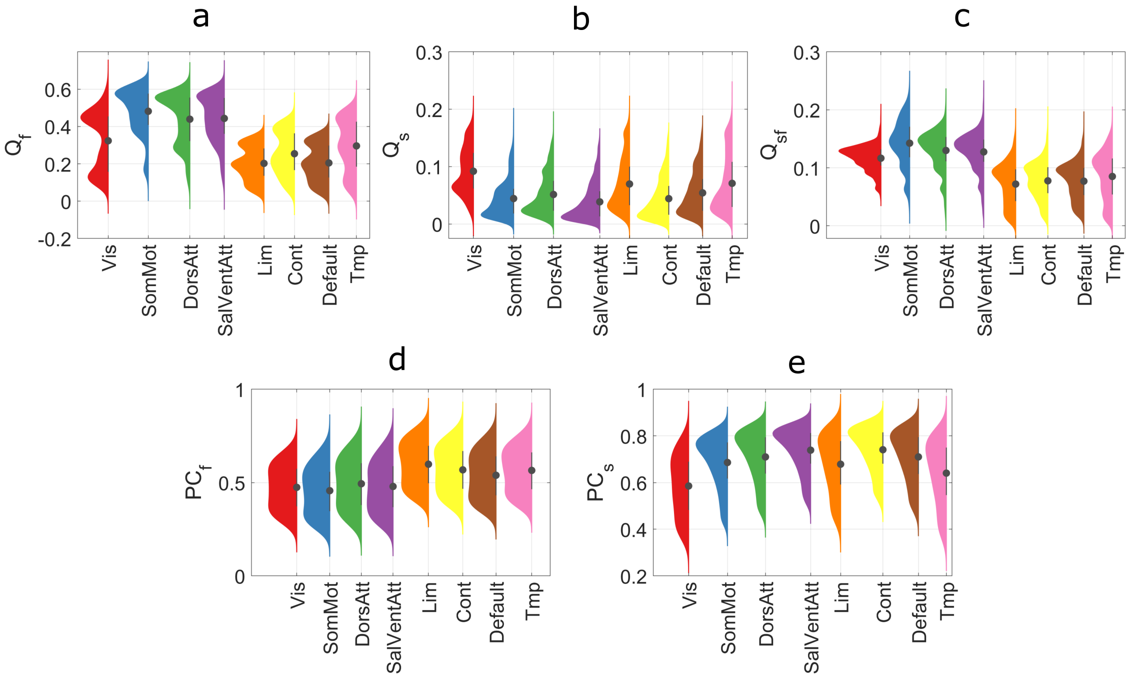


**Figure S12:** Top line: Contribution to functional (a), structural (b), and structure-function (c) modularity from single brain regions averaged across the Yeo canonical networks and across all combinations of $\{\gamma, \omega, \eta\}$. Bottom line: Participation Coefficient averaged across the Yeo canonical networks and all combinations of $\{\gamma, \eta\}$ when $\omega = 2.1210 - 6(\omega_{min})$.

### S8. Brain-behavior correlation

As a supplemental investigation, we propose a preliminary analysis of how patterns of variability between structural and functional community organization can be related to clinical assessments of the subjects. In addition to the imaging data, the NKI-RS projects also contains several clinical, behavioral, and cognitive measures from the participants, through a common protocol covering a wide array of psychiatric, cognitive and behavioral functions. Here we selected measures falling within the domain of the assessment of cognitive and executive functioning, the Wechsler Abbreviate Scale of Intelligence (WASI-II) (Wechsler, 1999) and the Wechsler Individual Achievement Test – Second Edition Abbreviated (WIAT-IIA) (Wechsler, 2005). The former scale (WASI-II) measures general intelligence, with IQ tests based on vocabulary, block design, similarities and matrix reasoning. It comprises the FSIQ (Full Scale IQ), an index taking into account all the sub-tests, the VIQ (Verbal IQ), which only considers the vocabulary, and the PIQ (Performance IQ), capturing the block design and matrix reasoning. The latter scale (WIAT-IIA) is an analogous set of tests to evaluate word reading, numerical operations and spelling. It results in a score that, from now on, we call COMP. Both tests can be administered to participants aged from 6 to 85 years old.

We investigated the correlation between those behavioral indices and the patterns of coupling between structural and functional partitions. In order to do that we extrapolated a subject-level measure associated to the patterns of SC-FC variability. The scheme in Figure S2e summarizes this procedure. We built a $N \times NS$ matrix, in which each column is referred to an individual, and is obtained by averaging the vectors of entropies computed between the SC-FC partitions laying in the parameters space where the principal components are most expressed (Figure 4a in the main article). Then, we calculated the Spearman correlation between each row of the matrix $(1 \times NS)$, indicating how much a node's assignment varies between SC and FC for each subject, and each one of the four cognitive assessment vectors (VIQ, FSIQ, PIQ, COMP).

We also investigated the correlation between the same clinical scales and the patterns of communities' variability across subjects.

Results are reported in Figure S13. For each component, we represented on the cortex surface the correlation coefficients statistically significant ($p < 0.05$) obtained considering for the VIQ index (figure S13(a-e)). These results suggested that while the entropy associated to some brain regions correlated similarly across different scales (identified by the different components), some other brain areas displayed a coupling between FC and SC partitions that correlates with VIQ in a different manner depending on the scale we are focusing on. The nodes involved in the DMN and visual area positively correlated with the coefficients of each principal component. This means that, in a subject, the more FC and SC partitions are uncoupled in these regions, the higher is the cognitive VIQ score achieved. The opposite happened in the nodes belonging to the temporal area, whose entropy negatively correlated with VIQ, meaning that the more the communities are matched between SC and FC in this region, the better results the cognitive assessment. The other brain regions present different trends of correlation passing from a subspace of the parameter space to another one. For instance, the entropy associated to the salience and ventral attention network positively correlated with VIQ in $PC_1$, $PC_2$, $PC_3$, $PC_4$, while negatively in $PC_5$. With respect to the others $PC_5$ is located at high $\gamma$ and low $\eta$ values, that is at combinations of the resolution parameters corresponding to a fine partition, mostly uncoupled between FC and SC networks. The somatomotor system was positively correlated with VIQ in the first two components, while negatively in the remaining three. Other systems showed a correlation only at certain scales, like the DAN, whose correlation coefficients in the $PC_1$ and $PC_4$ were close to zero, while in the $PC_2$ and $PC_3$ were positive and in the $PC_5$ negative.

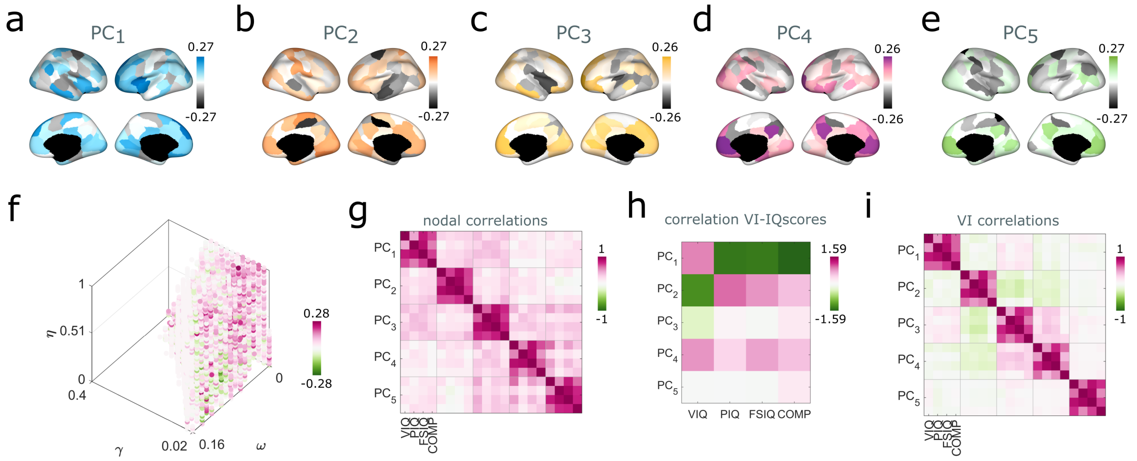


**Figure S13: Brain-behavior correlations.** (a-e) Projection on the cortex of the correlation coefficients obtained computing the Spearman correlation between each node's entropy and the verbal IQ index (VIQ), for the first five components. (f-g) Average within the functional systems of the correlation coefficients, for the five principal components. (k) Average of the correlation coefficients among nodes for each point in the parameter space. (l) Matrix encoding the values of the correlation coefficients obtained computing the Pearson correlation between node's entropy and IQ indices, for each pair of PC and IQ-index. (m) Matrix reporting the correlation coefficient of the Spearman correlation between the average VI of the partitions within each component (subjects' VI) and the four IQ indices. (n) Matrix encoding the values of the correlation coefficients obtained computing the Pearson correlation between subjects' VI and IQ indices, for each pair of PC and IQ-index.

In Figure S13(f) we showed the average values of the correlation coefficients for each point of the parameter space, in order to illustrate how this kind of brain-behavior correlations were highly sensitive to the resolution of the detected modules.

In Figure S14 we reported the results regarding the other three behavioral indices for cognitive assessment (COMP, FSIQ, PIQ). As shown by the matrix reporting the correlation between the correlations resulted from each pair of PC and behavioral index (Figure S13(g)), most of the variance in the correlation trends are observable $PC$-wise, while the trend across different behavioral indices are quite consistent.

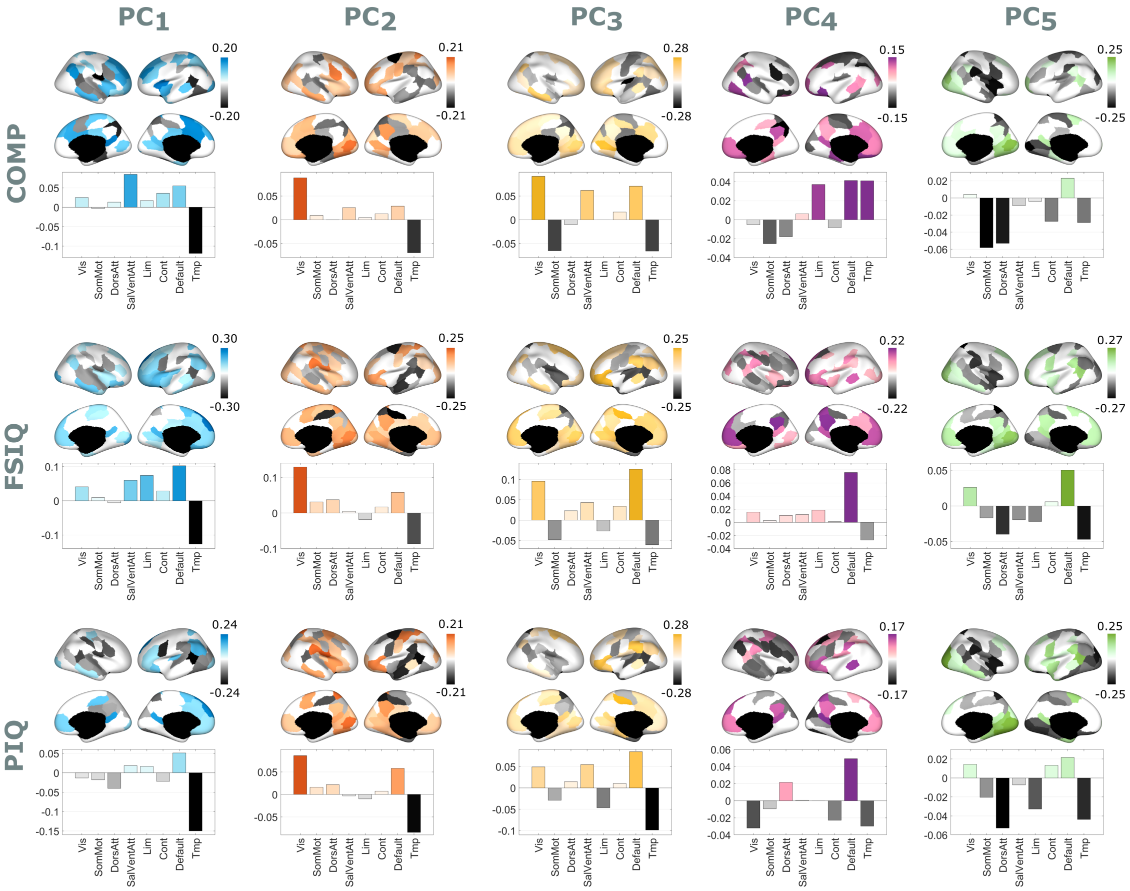


**Figure S14: Brain-behavior correlations.** Results of the same analysis shown for VIQ in Figure S13, replicated for the other three IQ indices: PIQ, FSIQ and COMP. Results are ordered so that each column is relative to one of the five principal components, while rows are relative to IQ indices. We show the projection on the cortex of the correlation coefficients obtained computing the Spearman correlation between each node's entropy and the IQ indices, together with the average within the functional systems of the correlation coefficients.

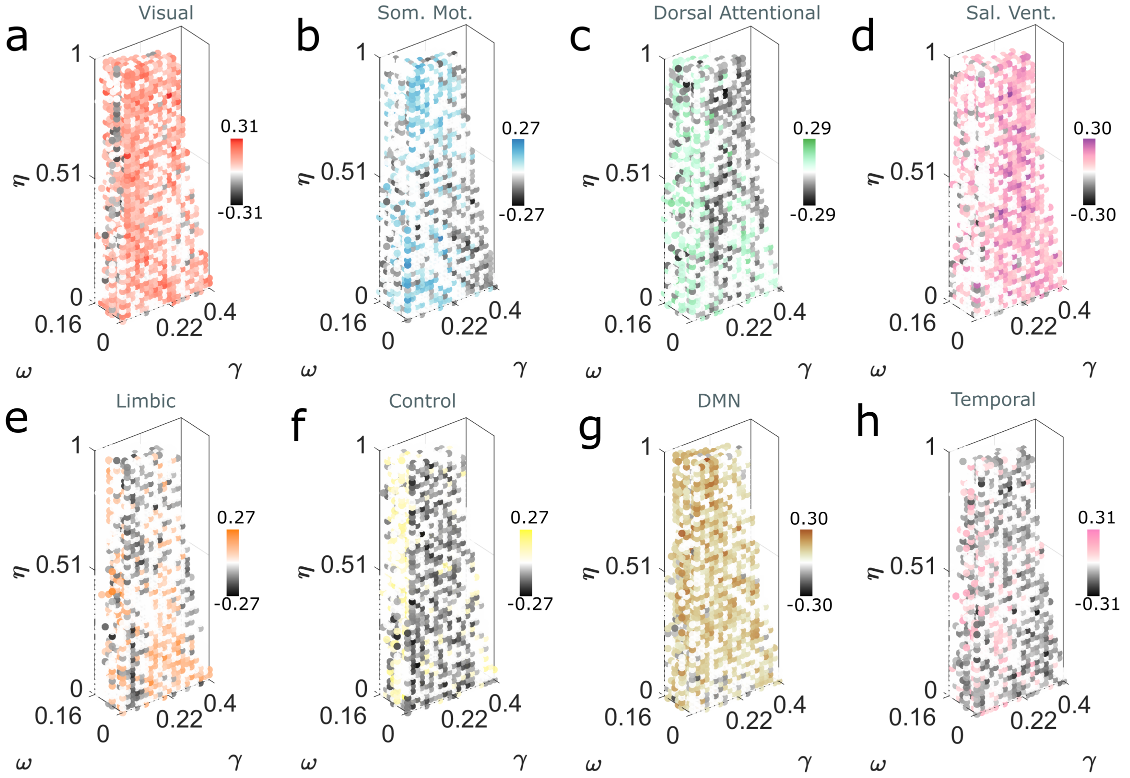


**Figure S15:** Average of the correlation coefficients within each functional system for each point in the parameter space. The correlation coefficients refer to the Spearman correlation computed between the community entropy (between SC and FC partitions) and the VIQ.

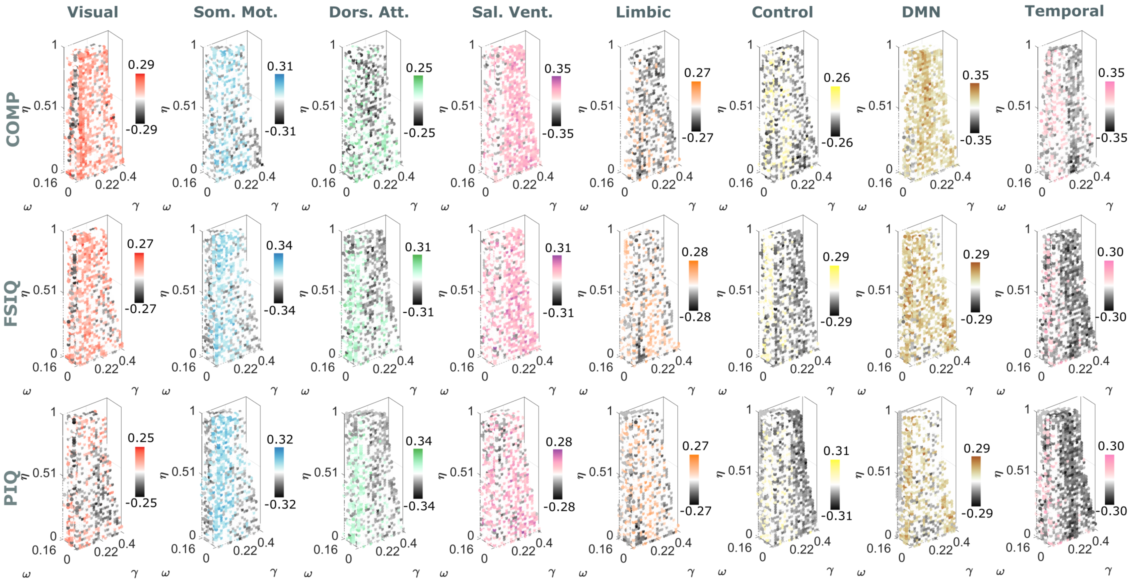


**Figure S16:** Average of the correlation coefficients within each functional system for each point in the parameter space. The correlation coefficients refer to the Spearman correlation computed between the community entropy (between SC and FC partitions) and the IQ indices named COMP, FSIQ and PIQ. Results are represented so that columns slide on functional systems, while rows slide on IQ indices.

### S9. Brain-behavior correlation considering the inter-subjects community entropy

We carried a similar analysis to investigate the relations between the IQ related cognitive indices (VIQ, COMP, FSIQ and PIQ) and the variability/coupling of the modular structure across subjects in both cases of functional and structural networks. For this purpose, we considered the first three components and we built two matrices (one for each type of connectivity) of dimension [nodes $\times$ subjects] in which each column is strictly related to the entropy between that subject's partition and the partitions of all the other subjects. Then, we computed the correlation between each row of the matrix and the behavioral indices.

Results are presented in Figure S17-S18. Also in this case the components are barely overlapped in the parameter space. The inter-subject modular variation of the regions involved in the temporal area is mainly positively correlated to clinical scale for the structural networks and negatively for the functional networks. As in the previous paragraph, the DMN presents an inter-subject variability that is always positively correlated to the considered behavioral scales, regardless of the partitions' resolution and the type of connectivity. Other areas instead, like the visual area or the DAN and ventral attention networks, present an inter-subject functional communities' variability which positively or negatively correlates with IQ according to the resolution at which we observe such communities. Patterns of correlation remain overall consistent across different IQ indices.

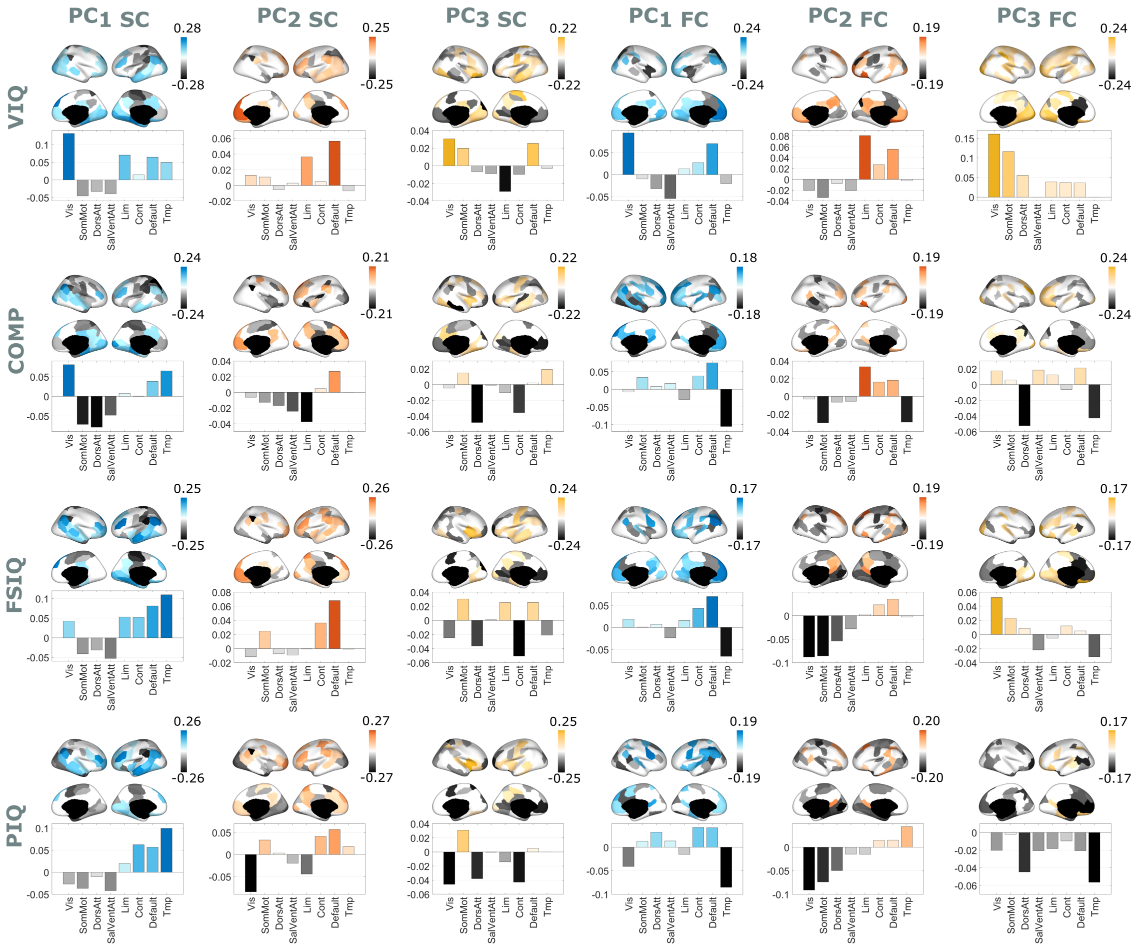


**Figure S17: Brain-behavior correlations within modality.** Results of the same analysis shown for the SC-FC analysis reported in Figure S13 and Figure S7, replicated for the inter-subject's analysis. Results are ordered so that the first three columns are relative to the three principal components of the analysis linked to structural networks, while the last three columns to the three principal components linked to the functional networks. The rows, instead, are relative to the four IQ indices, VIQ, COMP, FSIQ and PIQ. We show the projection on the cortex of the correlation coefficients obtained computing the Spearman correlation between each node's entropy and the IQ indices, together with the average within the functional systems of the correlation coefficients.

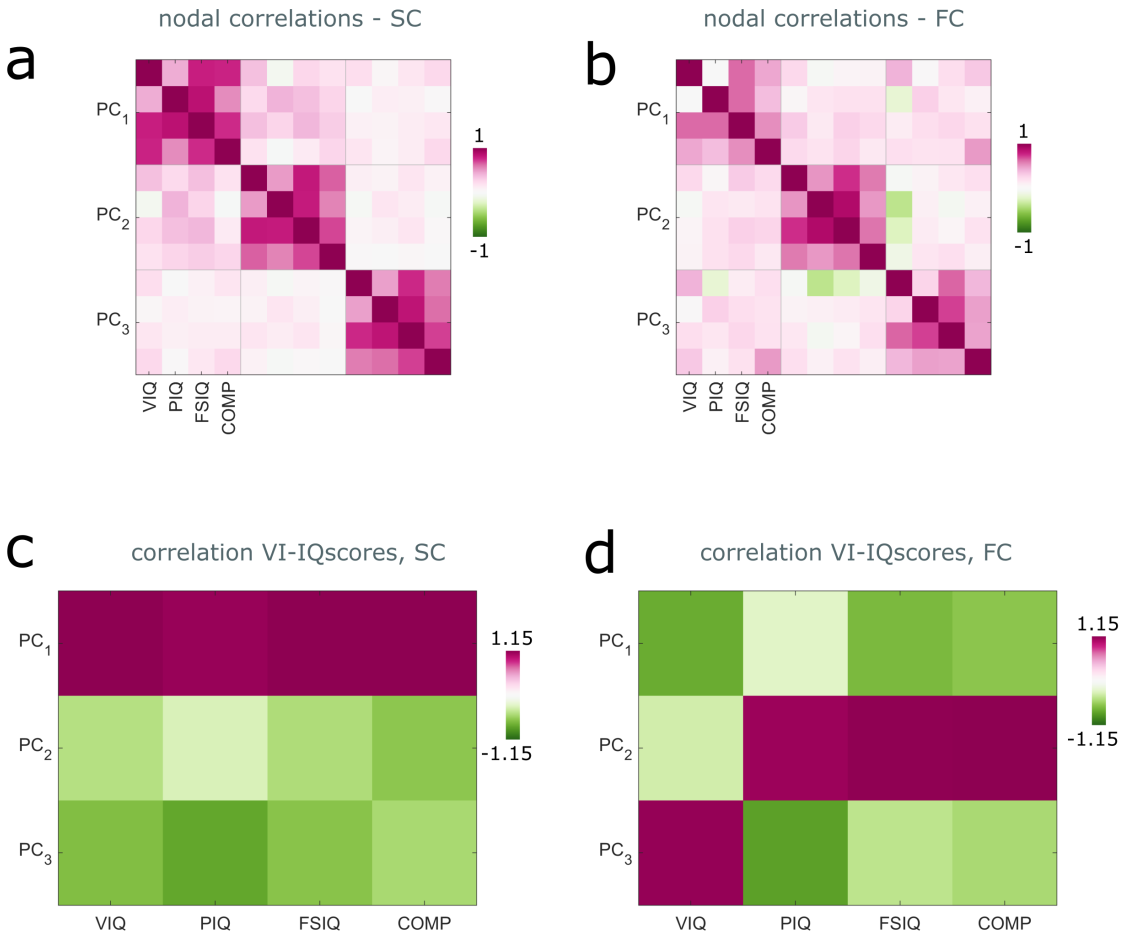


**Figure S18: Brain-behavior correlation within modality.** (a,b) Matrix encoding the values of the correlation coefficients obtained computing the Pearson correlation between node's entropy and IQ indices, for each pair of PC and IQ-index, for structural and functional connectivity networks, respectively. (m) Matrix reporting the correlation coefficient of the Spearman correlation between the average VI of the partitions within each component (subjects' VI) and the four IQ indices, for structural and functional connectivity networks respectively.
